# A versatile synaptotagmin-1 nanobody provides perturbation-free live synaptic imaging and low linkage-error in super-resolution microscopy

**DOI:** 10.1101/2023.01.30.525828

**Authors:** Karine Queiroz Zetune Villa Real, Nikolaos Mougios, Ronja Rehm, Shama Sograte-Idrissi, László Albert, Amir Mohammad Rahimi, Manuel Maidorn, Jannik Hentze, Markel Martínez-Carranza, Hassan Hosseini, Kim-Ann Saal, Nazar Oleksiievets, Matthias Prigge, Roman Tsukanov, Pål Stenmark, Eugenio F. Fornasiero, Felipe Opazo

**Author notes:** These Authors contributed equally to this work (Karine Queiroz Zetune Villa Real, Nikolaos Mougios, Ronja Rehm, Shama Sograte-Idrissi). This Author supervised the work (Felipe Opazo).

## Abstract

Imaging of living synapses has relied for over two decades on the overexpression of synaptic proteins fused to fluorescent reporters. This strategy changes the stoichiometry of synaptic components and ultimately affects synapse physiology. To overcome these limitations, here we introduce a nanobody that binds the calcium sensor synaptotagmin-1 (NbSyt1). This nanobody functions in living neurons as an intrabody (iNbSyt1) and is minimally invasive, leaving synaptic transmission almost unaffected, as demonstrated by the crystal structure of the NbSyt1 bound to synaptotagmin-1 and by our physiological data. Its single-domain nature enables the generation of protein-based fluorescent reporters, as we showcase here by measuring spatially-localized presynaptic Ca^2+^ with an NbSyt1-jGCaMP8 chimera. Moreover, its small size makes the NbSyt1 ideal for various super-resolution imaging methods. Overall, NbSyt1 is a versatile binder that will enable imaging in cellular and molecular neuroscience at a higher precision than possible in the past, over multiple spatiotemporal scales.

## Introduction

Neuronal communication strongly relies on the molecular machinery responsible for the accurate release of neurotransmitters. For this crucial step in neuronal communication, neurotransmitter-filled synaptic vesicles (SVs) fuse to the plasma membrane upon Ca^2+^ entry through activated voltage-gated channels following an action potential. The rapid fusion of the membranes from SVs to the plasma membrane occurs through a set of proteins known as the soluble N-ethylmaleimide-sensitive-factor attachment receptors or SNAREs. The SNAREs located at the plasma membrane form a four-helix bundle together with the SNAREs present at the SVs^1,2^. The interaction of these alpha-helices is progressive, and at some point, this complex provides sufficient energy for the fusion of the SVs with the plasma membrane, releasing the neurotransmitters’ load to the synaptic cleft. Interestingly, this SNARE-zippering mechanism for SVs fusion is regulated by Ca^2+^ in a complex manner. At central synapses, the putative protein responsible for the Ca^2+^-dependent regulation of synaptic release is Synaptotagmin 1 (Syt-1), which is present in several copies on each SV^3,4^. Syt-1 has two calcium-binding domains (C2A and C2B). Although the calcium dependency and Syt-1 involvement in evoked neurotransmitter release are generally not questioned, the exact molecular mechanism of how Syt-1 regulates the SNARE complex and SV fusion is a topic of fervent research^5–8^.

Remarkably, several questions remain to be answered, including which is the exact molecular target of Syt-1, what is the relative involvement of the two calcium binding domains, and ultimately which are the different steps and interactors of Syt-1. It is known for example that the C2 domains of Syt-1 interact with negatively charged membranes in a Ca^2+^-dependent manner^9,10^, although also a Ca^2+^-independent interaction has been observed in vitro for membranes containing phosphatidylinositol^11^. Several molecular mechanisms for Syt-1 have been proposed including the formation of rings on presynaptic membranes in the absence of Ca^2+^, which might prevent SNARE-mediated SV fusion^12^. One other model suggests that Syt-1 might stabilize a still yet little defined partially-assembled SNARE complex between the SV membrane and the plasma membrane and that this interaction with a partially-assembled complex is resolved in the presence of Ca^2+^, thus favoring fusion^13,14^. An additional model suggests that Syt-1 might specifically modulate the interaction with PIP2-containing membranes promoting changes in membrane bending in a Ca^2+^-dependent manner, thereby promoting fusion^9,15^.

We are persuaded that the heterogeneity of the experimental results obtained and the difficulty in understanding the molecular function of Syt-1 depend in part on diverging findings obtained *in vitro* and in cultured neurons^16^, and are in part related to the difficulty of studying this molecule when expressed at the correct stoichiometry^17,18^. For these reasons, here we aimed to develop a custom molecular tool that would allow us to manipulate and monitor the activity and the function of Syt-1 without adding a tag (e.g., EGFP) nor changing its stoichiometry in the cell, following a number of criteria.

We set out to develop a small probe able to bind to an accessible domain of Syt-1, with a strong affinity and high specificity, ideally, even if Syt-1 is engaged with the SNAREs, calcium or membranes in the crowded volume of an active synaptic bouton. The probe should be small to minimize the linkage error and retain its binding and specificity even after chemical fixation, which is typically needed for various advanced microscopy techniques, but it should also be functional in living neurons to monitor native Syt-1 location and function with minimal interference on Syt-1 physiological tasks.

We present here the successful development and characterization of an alpaca single-domain antibody, also known as nanobody (Nb), that remarkably fulfills most of our abovementioned criteria, and therefore it appears as an ideal probe to study living synapses and Syt-1 molecular physiology. This Nb is able to bind with high affinity and specificity to the C2A domain from rat Syt-1 and after obtaining the crystal structure of the Nb bound to rat C2A domain, we predicted that the Nb should significantly minimize the linkage error when directly conjugated to fluorophores and it should also impose a minimal effect on native Syt-1 activity. Due to these already favorable features, we decided to continue engineering this monovalent nanobody as an ideal probe to stoichiometrically reveal Syt-1 in various high- and super-resolution microscopy techniques like SIM, STED, DNA-PAINT, and Expansion Microscopy (ExM). In addition, we tested and demonstrated its minimal perturbation when expressing this Nb in living primary neurons (as intrabody), resulting in a flexible tool to label live synapses and follow synaptic vesicles. Finally, we used this specific binder to position the sensitive calcium sensor jGCaMP8^19^ directly on pre-synapses, allowing a highly localized calcium detection and fast response to evoked and spontaneous activity without genetic manipulation of synaptic proteins. We conclude that this Nb is a highly versatile tool for use in living neurophysiology studies as in conventional and super-resolution imaging techniques. We expect this Nb to be instrumental in tracking synaptic physiology and dissecting the molecular role of Syt-1 in neuronal communication.

## Results

### Selection and initial characterization of potential binders to Syt-1

For the selection of nanobodies against Syt-1 we immunized alpacas with enriched rat synaptosomes. Five months after the immunization protocol, we extracted the peripheral blood mononuclear cells (PBMCs) from 100 ml of blood. After total RNA extraction from the PBMCs we generated a cDNA library of Nbs and cloned them into a phagemid to later screen for binders using phage-display (Supp. Fig. 1a). Before performing the phage-display, we verified if these animals had generated heavy-chain antibodies against the cytosolic (amino acids 97-421) domain of rat Syt-1 (rSyt-1_(97-421)_). For this, we first removed conventional immunoglobulins (IgG1) that might also recognize this domain and enriched for heavy-chain antibodies (IgG2 & IgG3) from the plasma of the immunized alpacas (Supp. Fig. 1b). These IgG2 and IgG3 were used for the detection of rSyt-1_(97-421)_ coated as antigen in an ELISA plate (Supp. Fig. 1c). The ELISA result suggested that heavy chain antibodies binding rSyt-1_(97-421)_ were present in the serum although we used a complex antigen mixture for immunization (synaptosomes). These results encouraged us to proceed with phage-display screening.

After three rounds of phage display screening (biopanning) using the whole immobilized cytosolic rSyt-1_(97-421)_, we obtained five clones with strong positive signals on an ELISA. Although these clones have relatively similar complementary determining region 1 (CDR1), they all show significant differences in their CDR2s and CDR3s (Fig.1a, Supp. Fig. 1d). After subcloning all five clones into a mammalian expression vector, only two clones (A51 and A91) resulted in reasonable protein yields (>20 mg/L).

**Figure 1.**
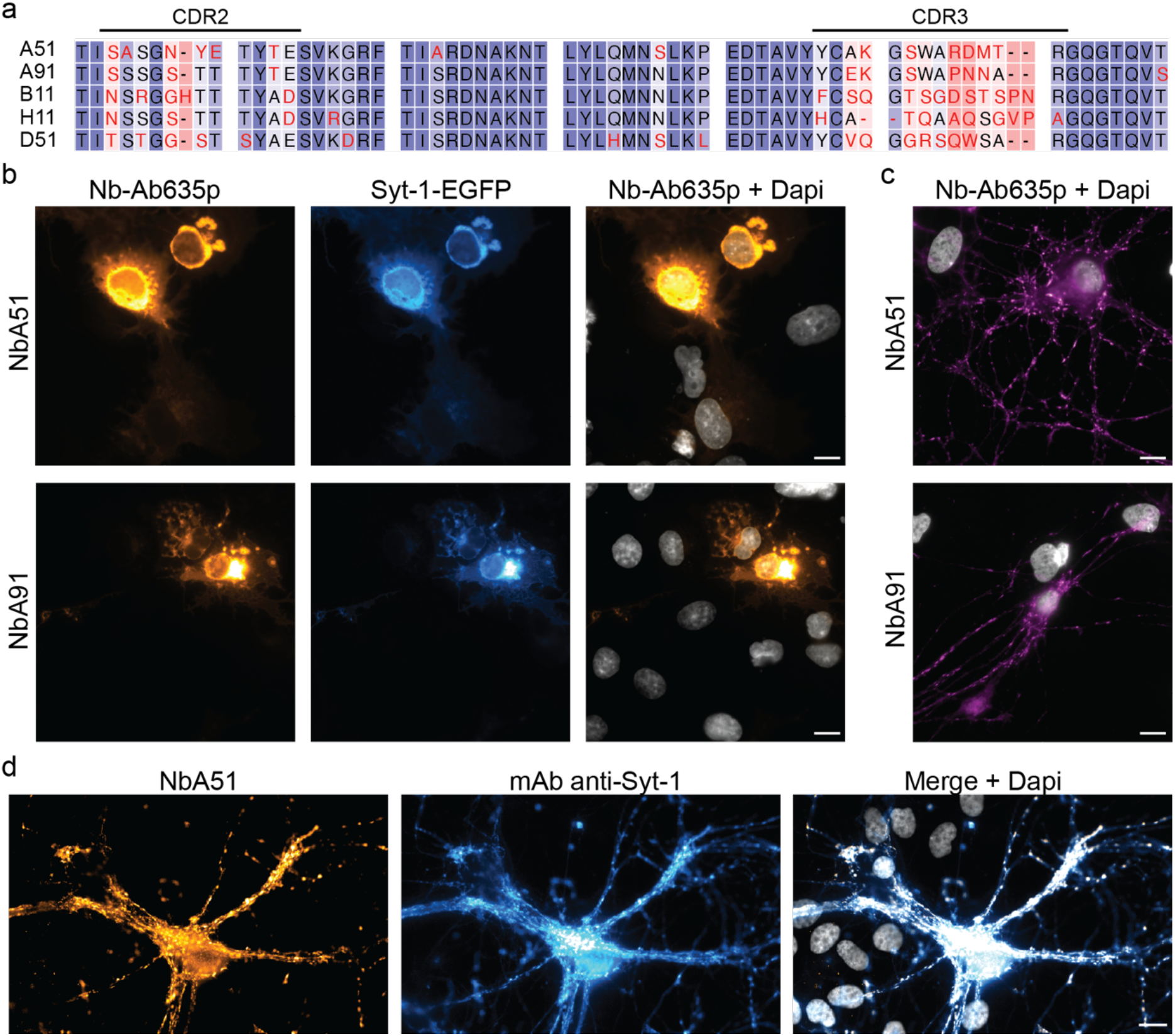
Selection of nanobodies against Synaptotagmin 1. (a) Alignment of five nanobody candidates selected for binding *in vitro* to rSyt-1_(97-421)_. Complementary determining regions (CDR) 2 and 3 show the maximum degree of variability. (**b**) Cos-7 cells transiently transfected with rat Syt-1 fused to EGFP (Syt1-EGFP) stained with NbA51 and NbA91 directly labeled with AbberiorStar635p (Nb-Ab635p). Scale bar 10 μm. (**c**) Hippocampal primary culture stained with Nb-Ab635p. Scale bar 10 μm. (**d**) Co staining of primary neuronal culture using the NbA51-Ab635p and a conventional monoclonal antibody revealed by polyclonal secondary antibody conjugated to Cy3. Scale bar 10 μm. Nuclei are stained with DAPI and depicted in greyscale.

For further validation, NbA51 and NbA91 were directly conjugated with fluorophores for testing them in immunofluorescence (IF) assays. Initially, COS-7 fibroblasts were transfected with plasmids coding for the full-length rat Syt-1 fused to EGFP. After 24h, cells were chemically fixed using paraformaldehyde and IF was performed using 50 nM of fluorescent nanobodies. Both clones displayed specific signals that were detected only on EGFP-positive cells, suggesting that both clones are able to detect rat Syt-1 and that their epitopes seem not to be affected by aldehyde fixation (Fig. 1b). In order to understand the specificity of these candidates, a dot-blot assay with purified rSyt-1_(97-421)_ and rSyt-2_(97-422)_ suggest that both candidates bind specifically to rSyt-1 and not rSyt-2 (Supp. Fig. 2). We carried on an additional validation step to verify their IF performance on rat primary hippocampal cultures. At this point we reasoned that if the nanobody would bind efficiently in a cell context situation, where Syt-1 is in close proximity with several other interacting molecules, it would thus be the one providing the higher signal-to-noise ratio (SNR) likely because has a higher affinity or is able to find more epitopes. While both candidates tested displayed the classical punctuated pattern of synapses expected in these primary neurons, by closer inspection the candidate NbA51 provided images with negligible level of background staining in cell bodies of neurons and astrocytes. Due to this, we decided to continue to work with the NbA51 and rename it NbSyt1.

To further confirm the specificity of NbSyt1, we performed the co-staining of primary hippocampal neurons using a directly labeled version of the nanobody and a conventional monoclonal antibody anti-Syt-1, which displayed virtually a complete colocalization on confocal microscopy (Fig. 1d), confirming that this NbSyt1 appears to be also specific in a neuronal environment. Next, we decided to understand more precisely where the NbSyt1 binds within rSyt-1_(97-421)_. For this, we produced various fragments of rSyt-1_(97-421)_ and performed a dot-blot assay. The analysis of these experiments indicated that the NbSyt1 binds to the C2A calcium-binding domain of Syt-1. However, if the putative protein linker between C2A and C2B is removed (fragment-4 & fragment-6, Supp. Fig. 2b) the affinity was strongly reduced or completely lost. This finding motivated us to proceed with the crystallization of the complex NbSyt1-Syt-1 and to resolve their interaction at high resolution.

### The structure and interaction affinity of the complex NbSyt1 and C2A domain of rSyt-1

In order to perform X-ray crystallography structural studies, the NbSyt1 was expressed and purified in its minimal size and without tags to be mixed in equal molar ratio with the tag-free rSyt-1 C2A_(140-265)_ domain. A crystal diffracting to 1.68 Å revealed the NbSyt1: C2A_(140-265)_ complex (PDB 8C5H; Supp. Table 1), burying a contact interface of approx. 662 Å^2^. Both proteins have similar folds, consisting of two antiparallel beta sheets and short alpha-helical segments. Most contacts are between side chains on both the NbSyt1 and C2A_(140-265)_, except for S107 on the NbSyt1 and Q154 and Q209 on C2A_(140-265)_, which make contacts with the backbone of the neighboring chain. As shown in Figure 2d, F40 from the NbSyt1 extends into a hydrophobic pocket on C2A_(140-265)_ and is sandwiched between N207 and Q209. The C2A_(140-265)_ aspartate residues 172, 178 and 230 coordinate a calcium ion, and the presence of the NbSyt1 does not seem to affect the calcium binding of the C2A domain. The residue E257 on the crystal structure on the C2A_(140-265)_ has contact with 2 amino acids on the NbSyt1. Interestingly, E257 was missing in fragment-4 when performing the epitope mapping (Supp. Fig. 2b), which could explain the lack of signal of NbSyt1 on fragment-4, but a strong signal with fragment-6 (including E257). Next, we took the full cytosolic domain rSyt-1_(97-421)_ and determined with the microscale thermophoresis assay that the NbSyt1 binds with an affinity of 0.7±0.3 nM to rSyt-1_(97-421)_ (Supp. Fig. 3).

**Figure 2.**
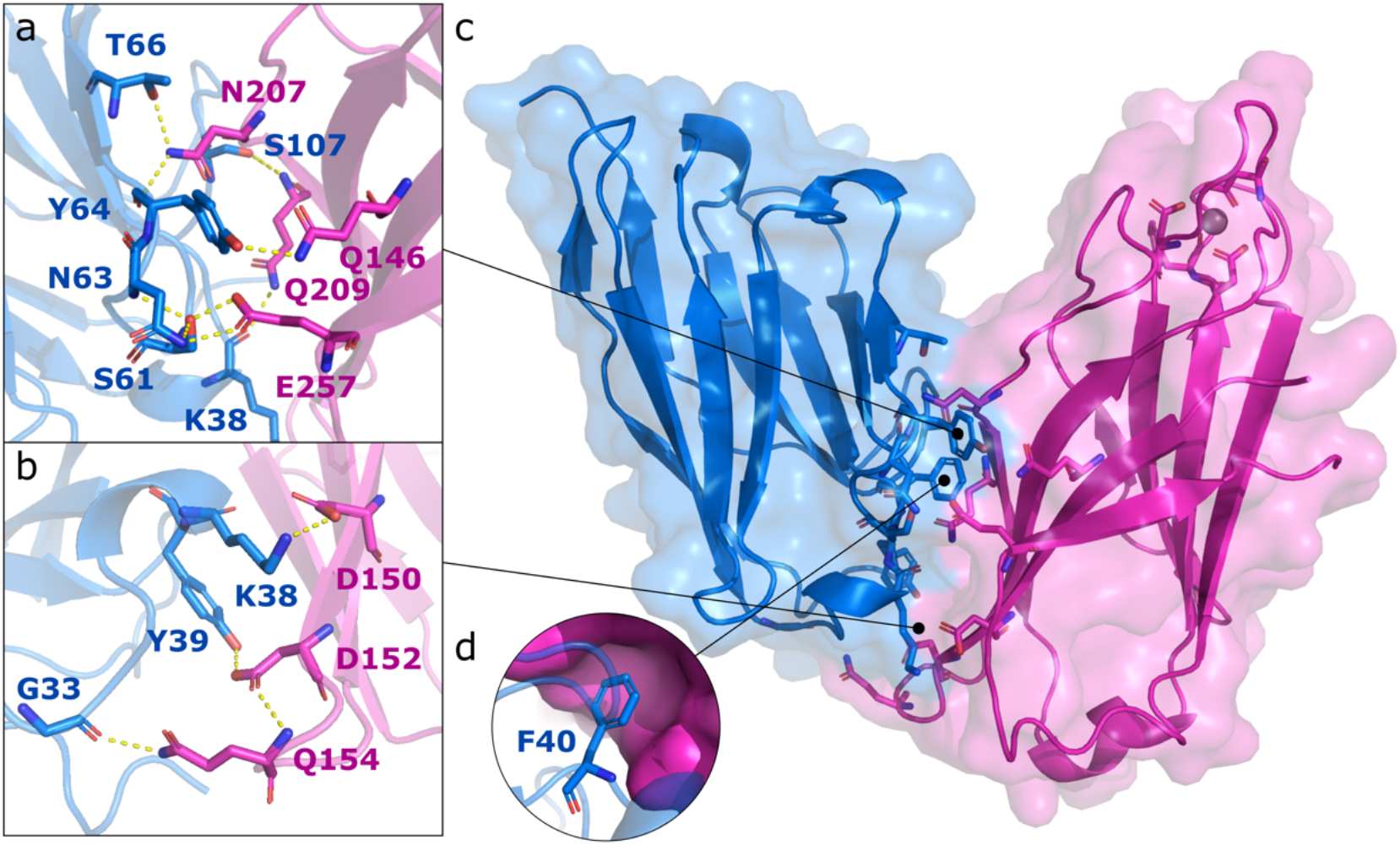
NbSyt1 (blue) in complex with its target rat Synaptotagmin 1 cytosolic C2A_(140-265)_ domain (magenta), PDB ID: 8C5H. (**a**, **b**) Polar contacts between amino acids of the NbSyt1(blue) and C2A_(140-265)_ domain (pink), (**C**) Overall structure of the complex, (**d**) Hydrophobic interaction between F40 in the NbSyt1 and a hydrophobic pocket in Syt-1 C2A_(140-265)_ domain.

When analyzing the structure to understand why the NbSyt1 interacts very weakly with rSyt-2, two points were observed. First, the glutamine Q209 in rSyt-1 is exchanged for threonine in rSyt-2. This key interacting residue has direct contact with the backbone of K38 and F40 and a polar contact with side chain of residue S107 on the NbSyt1. Secondly, the valine V255 in rSyt-1 is exchanged for proline in rSyt-2, which is expected to alter the backbone geometry of the polypeptide chain and disrupting interactions in its proximity, e.g. the interactions between E257 (conserved in rSyt-2) and the NbSyt1 (S61 & N63).

### Revealing Syt-1 with minimal linkage error in various super-resolution microscopy techniques

After determining the NbSyt1 binding affinity, binding epitope and specificity to Syt-1, we conjugated the NbSyt1 directly to fluorophores or to single-stranded DNA (ssDNA), to fulfill some of the requirements for various advanced imaging techniques like Stimulated Emission Depletion (STED), Points Accumulation for Imaging in Nanoscale Topography (PAINT) and Structural Illumination microscopy (SIM). Based on the structure, we know that the linkage error from the fluorophore on the N-terminus of the NbSyt1 is only ~1.2 nm to the closest part of the C2A, while the fluorophore located on the C-terminus is slightly below 4 nm from its target. In the case of DNA-PAINT imaging, the Atto488 fluorophore on the imager strand (i.e., the ssDNA annealing to the conjugated ssDNA on the NbSyt1) will be approximately ~4 nm away from Syt-1. This means that for the super-resolution imaging techniques tested here, the small linkage error imposed by the NbSyt1 is neglectable. Results shown on Figure 3 suggest that the commercial STED and SIM setups provided a slightly lower resolution as compared to single-molecule localization-based techniques: Exchange-PAINT ^20,21^ and Fluorescence Lifetime-PAINT (FL-PAINT) ^22^ strategies. This becomes evident when looking into the gaps in intensity profiles between the Syt-1 localization peak (at the pre-synapse) with the postsynaptic density marker PSD-95 peak, also detected with a directly labeled nanobody (Fig. 3bc). Exchange-PAINT had the tendency, in our comparison, to provide a more accurate distance between pre- (Syt-1) and post- (PSD-95) synaptic compartments (i.e., ~100-150 nm^23,24^).

**Figure 3.**
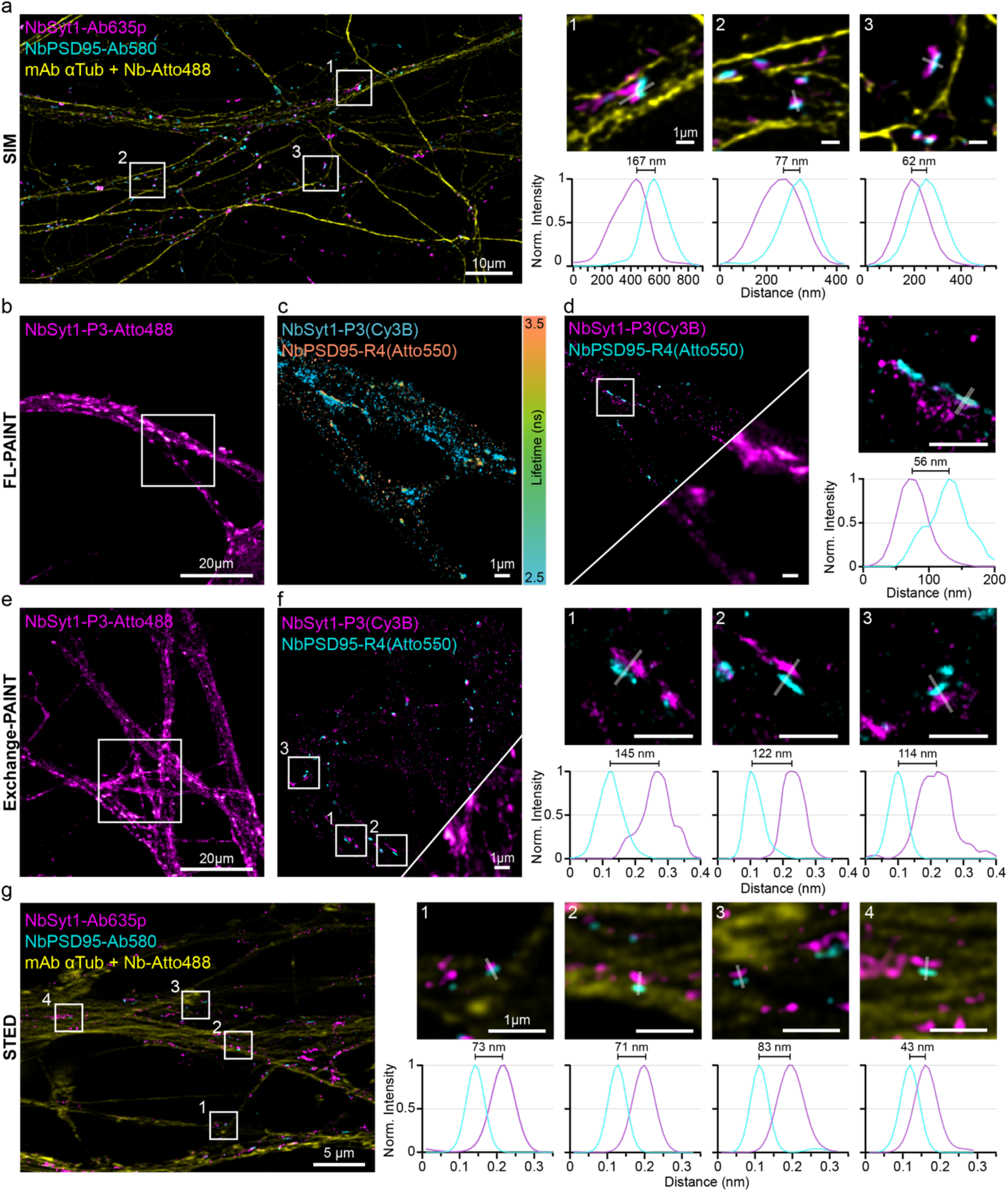
Imaging of NbSyt1 in various advanced imaging techniques. Primary hippocampal neurons were stained and imaged. **(a)** Full field of view from Structured Illumination Microscopy (SIM) of NbSyt1 (magenta), NbPSD95 (cyan) and mAb anti-αTub (yellow). Nbs were directly conjugated with the fluorophores and mAb anti-αTub was revealed with anti-mouse IgG1 secondary nanobody (2.Nb-Atto488). White squares denote the magnified regions used for the normalized fluorescent intensity line profiles of NbSyt1 (magenta) and NbPSD95 (cyan). **(b)** Diffraction-limited imaging of neurons stained with NbSyt1-P3-Atto488. White square indicates the region displaying fluorescence lifetime PAINT (FL-PAINT). (**c**) Fluorescence lifetime imaging of NbPSD95-R4 and NbSyt1-P3-Atto488 with the P3-Cy3B and R4-Atto550 imager strands. Image is color coded based on the recorded lifetimes from 2.5 to 3.5 ns (see methods and Supp. Fig. 4). (**d**) Result image of NbSyt1-P3-Atto488 and NbPSD95-R4 after lifetime segmentation. The white square indicates the zoomed regions with a the normalized fluorescent intensity line profiles of NbSyt1 (magenta) and NbPSD95 (cyan). **(e)** Diffraction-limited imaging of NbSyt1-P3-Atto488. White square indicates the region selected to display (**f**) Exchange PAINT imaging of NbSyt1-P3-Atto488 (magenta) and NbPSD95-R4 (cyan) using P3-Cy3B and R4-Atto550 as imager strands. The zoomed-in regions are indicated by the white squares 1, 2 and 3 with the normalized fluorescent intensity line profiles of NbSyt1 (magenta) and NbPSD95 (cyan). **(g)** Stimulated Emission Depletion (STED) imaging of NbSyt1 (magenta), NbPSD95 (cyan) and mAb anti-αTub (yellow). Nbs were directly conjugated with fluorophores and mAb anti-αTub was revealed with 2.Nb-Atto488. White squares show the chosen regions of interest for the normalized fluorescent intensity line profiles of NbSyt1 (magenta) and NbPSD95 (cyan). All distanced denoted between the Syt-1 and PSD-95 signals are calculated from the max value after Gaussian fit of the intensity line profile.

Next, we tested if this nanobody could be used in expansion microscopy (ExM). It has been suggested that nanobodies might not work in all ExM techniques^25^. Maybe due to their small size and limited numbers of fluorophores, the signal from nanobodies might not be well retained in some of the hydrogels used (Fig. 4a). Although this can happen, it strongly depends on the actual sequence of the Nb and the ExM technique. Some nanobodies could be more efficiently retained than others. With this limitation in mind, we modified the NbSyt1 by adding 4 extra lysines flanking the ectopic cysteines on their N and C-termini where the fluorophores are site specific conjugated (ExM cassettes). The sequence is not digested by Proteinase K (used for some ExM techniques) and added residues with negative charges to compensate the positive charges introduced by the lysines. We aimed that these cassettes should help retaining the fluorophores to the gel following fixation for ExM. Eventually, we managed to perform 10X ExM^26^ with directly labeled NbSyt1 bearing two expansion microscopy cassettes (ExNbSyt1) and we imaged Syt-1 on expanded synapses together PSD-95 revealed by co-staining with a nanobody anti-PSD-95 (Fig. 4d).

**Figure 4.**
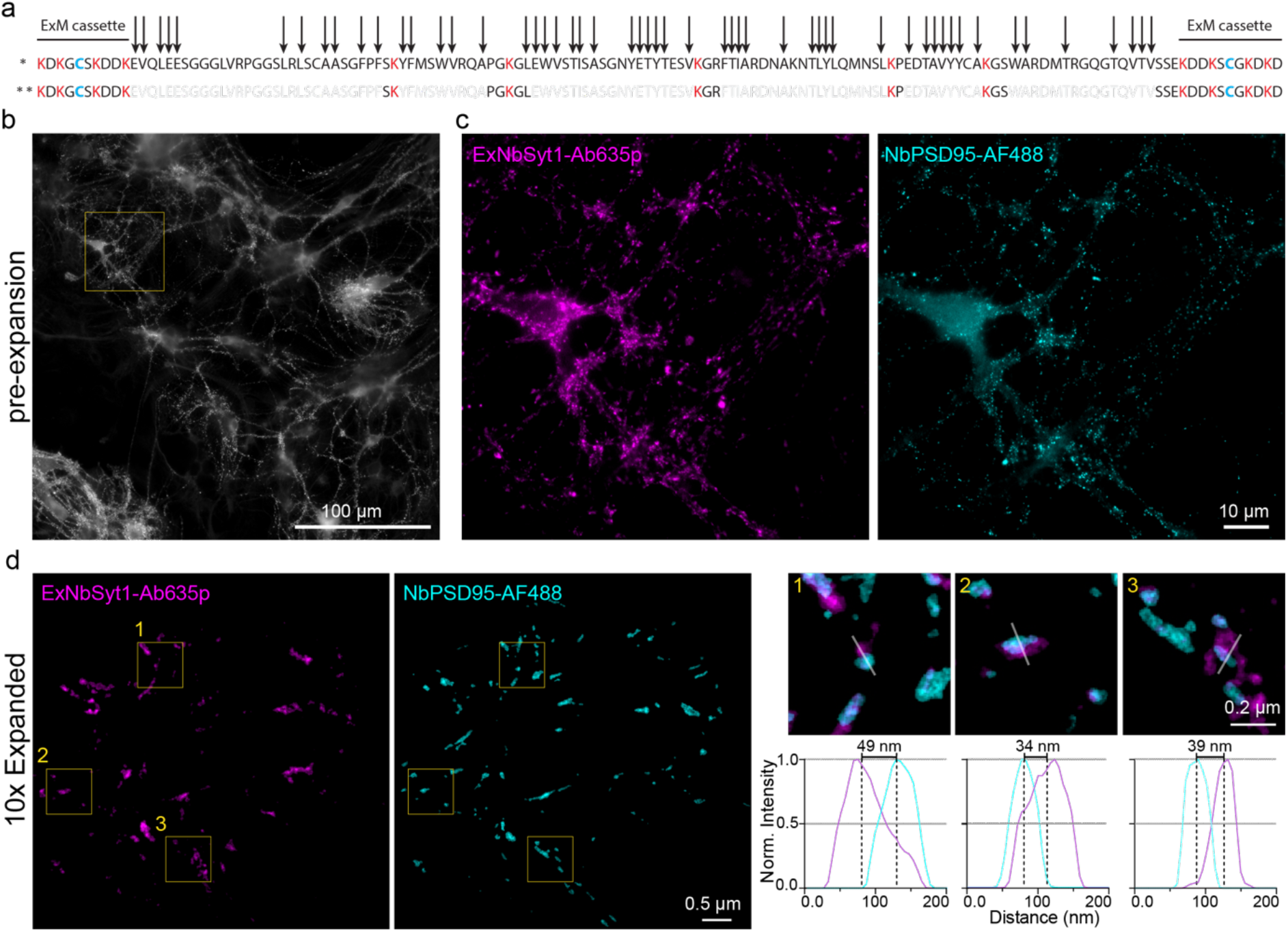
Tailored NbSyt1 for post-fixation or anchoring to ExM hydrogels. (**a**)We modified the NbSyt1 by adding ExM cassettes on its N and C-termini. This introduced four lysines on each end to ensure that fluorophore gets anchored on the gel. Arrows represent theoretical cleavage sites by Proteinase K, and ** displays the fragments that could be retained on a gel after complete proteinase digestion. (**b**) Epifluorescence large field of view of cultured neurons-stained pre-expansions with ExNbSyt1-Ab635p. (**c**) Magnified region on b displaying ExNbSyt1-Ab635p (magenta) and FluoTag-X2 anti-PSD95 AlexaFluor488 (cyan). (**d**) The same sample as in b is now expanded and imaged in a laser-scanning confocal microscope. Squares show chosen regions of interest and the corresponding normalized fluorescent intensity line profiles of ExNbSyt1 (magenta) and NbPSD95 (cyan).

### Using NbSyt1 as intrabody (iNbSyt1) in living neurons

The single-chain nature of nanobodies, allows their expression inside eukaryotic cells. Nanobody-based intrabodies^27^ have been used to follow target protein in living cells and create molecular sensors for in vivo applications ^28^. To test if the NbSyt1 folds properly in the reducing cytoplasm of living neurons and retains its high epitope specificity towards Syt-1, we subcloned the NbSyt1 into an adeno associated virus (AAV) vector in frame with its C-terminus to mScarlet^29^ or mNeonGreen^30^ fluorescent proteins. Primary hippocampal neurons tolerated the infection and expression of this chimera for more than 4 days without showing any obvious signal of toxicity (Supp. Movie1 & Supp. Movie2 for mScarlet and mNeonGreen respectively). Moreover, when imaged live, we were able to follow packages or synaptic vesicles moving along neuronal processes, providing a high contrast and good signal to noise ratio (Fig. 5 & Supp. Movie 1). This provided us the initial indications that the NbSyt1 is functional as an intrabody (iNbSyt1) in primary hippocampal cultures.

**Figure 5.**
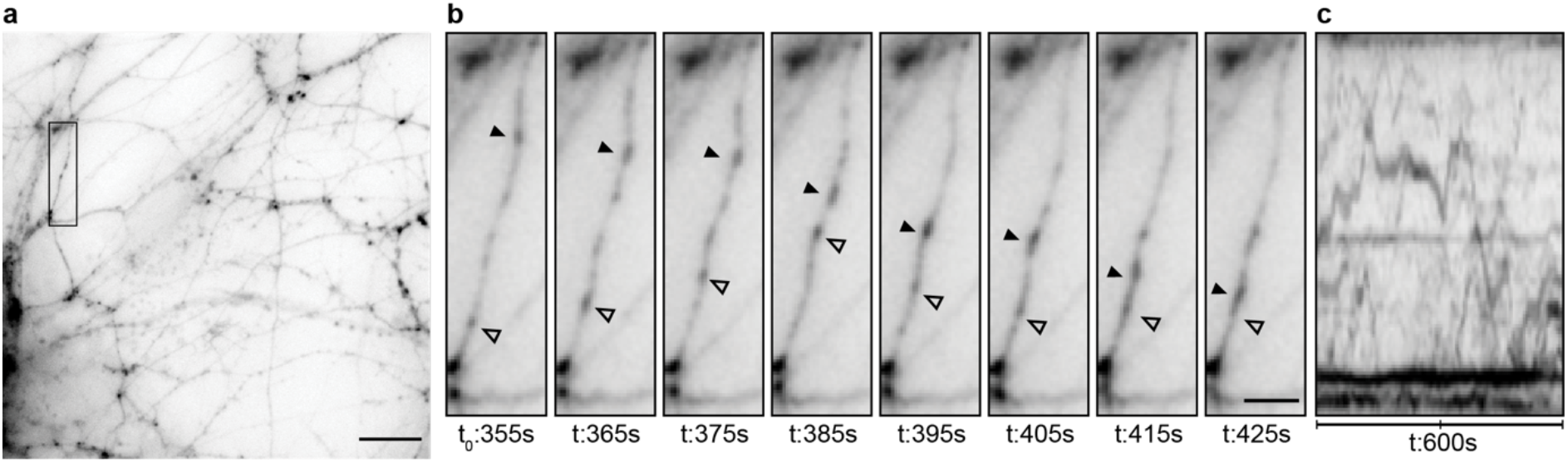
Live imaging of iNbSyt1-mScarlet. (**a**) Full field of view of a timelapse imaging (Supp. Movie 1). Area inside the black rectangle is magnified in b. Scale bar represents 10 μm. (**b**) Crop region of time lapse imaging at different timepoints (t). Arrow heads denote iNbSyt1mScarlet signal moving along a neuronal process. Scale bar represents 5 μm. (**c**) Ten minutes kymograph of the neuronal process.

### Assessment of synaptic activity in presence of the iNbSyt1

To test whether our engineered NbSyt1 intrabody with its high specificity to the C2A domain of Syt-1 is not altering synaptic physiology, we investigated neuronal activity in primary neuronal culture. We first checked changes in synaptic vesicle release properties using the fusion of VAMP2 with the superecliptic GFP, also known as SynaptopHluorin (SpH)^31^. This approach allowed us to evaluate if the iNbSyt1 has any effect on synaptic vesicle release and SV endocytosis. First, we confirmed that the iNbSyt1 is clearly localized at presynaptic terminals where it strongly colocalized with immunostainings for Synaptophysin and Bassoon (Fig. 6a). Then, we generated a bicistronic AAV expressing the iNbSyt1 and SpH^31^. When following the SpH signal after 60 action potentials (AP) or 600 AP at 20 Hz stimulation rate, we were not able to observe differences in SyH signal traces between neurons with or without expression of the iNbSyt1 (Fig. 6b). These results directly suggest that the expected enrichment of iNbSyt1 at pre-synapses, does not affect synaptic vesicle fusion mechanisms under evoked release, nor the endocytosis and subsequent reacidification of vesicles.

**Figure 6.**
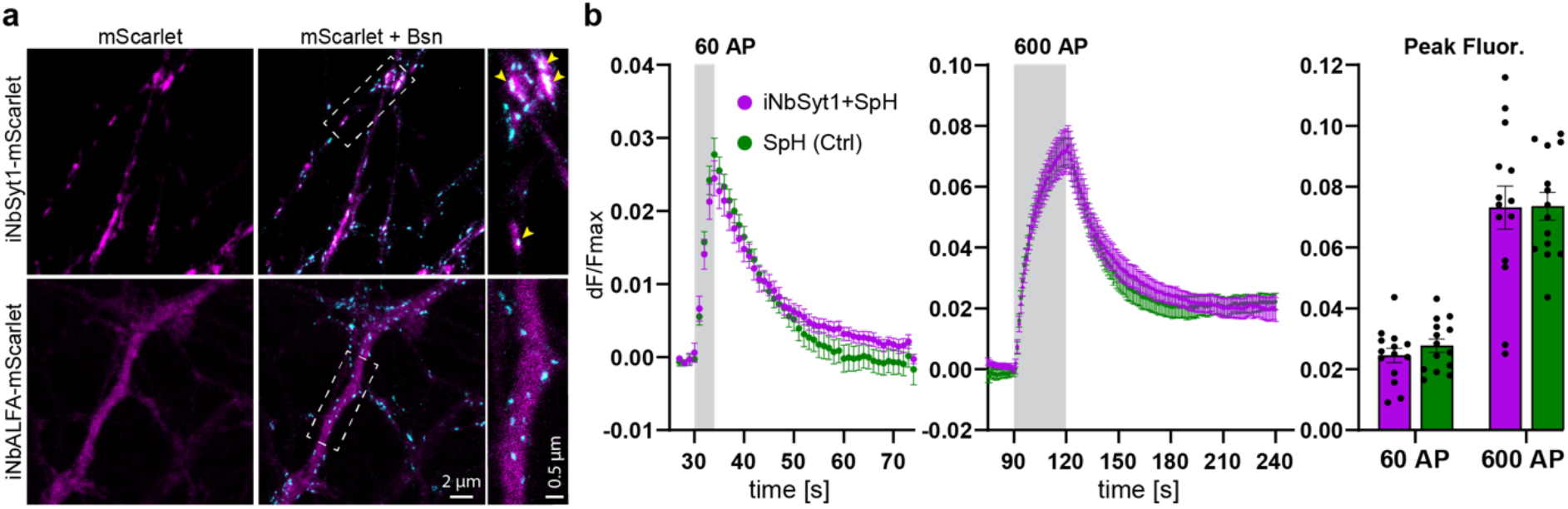
Intrabody iNbSyt1 localization and its potential effects on exocytosis physiology monitored using SynaptopHluorin. (**a**) The NbSyt1 as intrabody fused to mScarlet (iNbSyt1-mScarlet) displayed in magenta gets enriched in synapses confirmed by its co-localzation signal (arrow heads on zoomed region) of immunostained Bassoon (Bsn) imaged under STED microscopy showed in cyan. Expression of the NbALFA fused to mScarlet as intrabody (iNbALFA-mScarlet) provides a diffused signal distribution throughout neuronal processes. Scale bar represents 2 μm and 0.5 μm in zoomed region. (**b**) Neurons AAV infected with a bicistronic expression of iNbSyt1 and SynaptopHluorin (SpH) enabled to monitor the iNbSyt1 potential effect on synaptic vesicle fusion competency and dynamics. The expression of only SpH as control, resulted in no significant difference in peak fluorescence nor in rise or decay time of pHluorin signal after electric field stimulation with 60 action potentials (AP) or 600 AP at 20 Hz. Plots in (**b**) display the average and the SEM from 14 independent experiments (n=14), approximately 200 ROIs automatically were selected and analyzed per experiment with a total of 3323 ROIs for iNbSyt1 and 2860 for NbALFA.

Next, we looked for changes in the spontaneous release rate that could potentially arise from expression of our Syt-1 intrabodies. We therefore recorded non-evoked excitatory and inhibitory postsynaptic currents in the presence of tetrodotoxins in primary neuronal culture (miniature EPSCs and IPSCs) (Fig. 7a) for both, the iNbSyt1 and the original candidate iNbA91. We observed no significant differences in the amplitudes (quantal release of neurotransmitters) nor in Tau_(on)_ and Tau_(off)_ kinetics between cultures infected with iNbSyt1 and iNbALFA^32^ (used as a control intrabody since it has no target molecules in neurons). For the mEPSC rate, we observed a trend for both iNbSyt1 (iNbA51) and iNbA91 with a slight increase in rate of miniature postsynaptic currents, however not statistically significant for iNbSyt1 (Fig. 7d).

**Figure 7.**
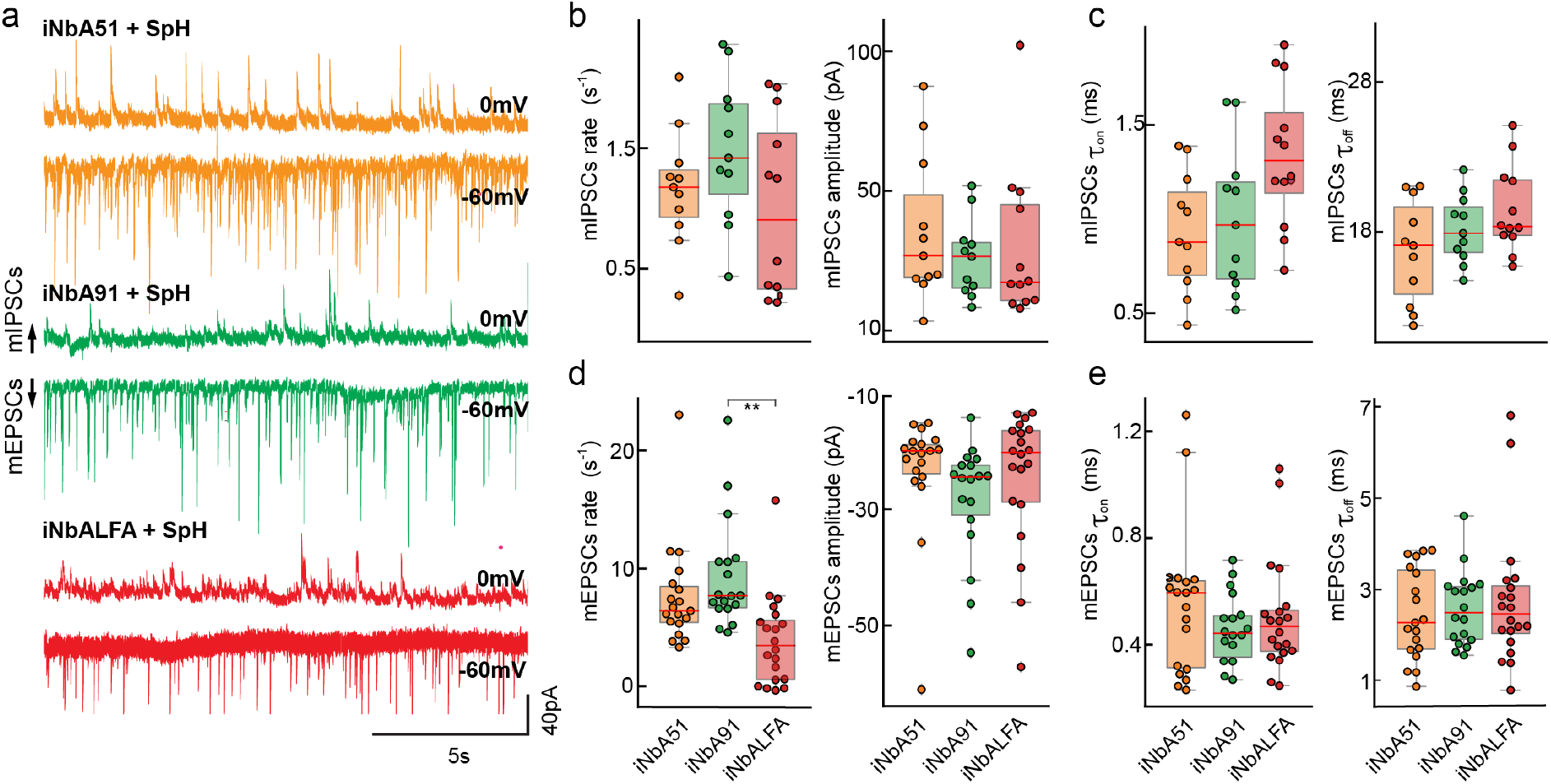
Electrophysiological analysis of mini EPSCs and IPSCs on primary hippocampal neurons. (**a**) Exemplary traces of mEPSCs and mIPSCs of neurons in a whole-cell configuration clamped at −60 and 0 mV, respectively. Cultures were infected using AAVs containing sequences of the respective intrabody and SpH. Upper traces represent mIPSCs and lower traces mEPSPs. **(b-d)** Boxplots summarizing mIPSCs and mEPSCs properties such as rate and amplitude (b and d) and Tau(on) and Tau(off) (c and e). For statistics, mIPSCs, n = 10-12 cells per group; mEPSCs, n = 18-21 cells per group, a one-way ANOVA with multiple comparison was applied with Tukey correction. *P*-values: ** = *p* < 0.01.

In light of our structural data that shows binding of NbSyt1 to the C2A domain of Syt-1, these findings are in line with the observations made by Courtney et al.^8^. They also observed that after genetically removing the C2A domain from Syt-1, a minimal effect on evoked release and a slight increase in the spontaneous miniature release could be observed. On the contrary, they also show that if the C2B domain is truncated, the action potential triggered synaptic vesicle fusion to the plasma membrane is drastically impaired^8^. Therefore, our data suggest that the iNbSyt1 bound to the C2A has no detrimental physiological effect in evoked activity on the synaptic physiology, and potentially neglectable effects in facilitating spontaneous release.

### Targeted calcium detection on pre-synapses

After characterizing the nanobody, its interactions and its effects on synaptic physiology, we thought to use it as an ideal tool to increase the synaptic targeting of genetically encoded calcium indicator, to be able to measure the Ca^2+^ changes in close proximity of the SV sensor for Ca^2+^ (i.e., Syt-1). This approach would allow us to reveal active synapses in living neurons, and perform calcium imaging on pre-synapses with high precision and with minimal impact on neuronal physiology, as we are not overexpressing chimeric synaptic vesicle proteins for the targeting. For this we generated new AAVs expressing the NbSyt1 as an intrabody and fused to the calcium sensor jGCaMP8s and jGCaMP8m (iNbSyt1-jGCaMP8)^19^. As control, we used the iNbALFA fused to the calcium sensors. As the NbALFA has been shown to work as intrabody^32^, but has no specific binding partners in rat hippocampal neurons, this would be the most appropriate control for these experiments. As expected from previous results, we observed the enrichment of synaptically-targeted calcium sensors enriched at synaptic boutons (Supp. Movie 3), and mostly a diffused signal with the control (Fig. 8a). Both sensors showed good signal following evoked 1 and 3 APs (Fig. 8b). However, when observing the traces from single boutons, the synaptically localized sensor provided more reliable signals, as single action potentials were sometimes lost in the background when measuring with the control sensor (Fig. 8b). To further corroborate this observation, we performed a more formal analysis on several boutons, also considering if the synaptically targeted sensor provides additional spatial information for the calcium signal compared to the control. For this we identified the boutons at the end of each movie following a 300 AP stimulus, and used this signal to select the center of the boutons synaptic proximal and distal areas (Fig. 8c). Using this analysis, we were able to see that the pre-synaptically localized sensors provided more precise spatial resolution and better signal to noise ratio (SNR) when compared to the control (Respectively Fig. 8d, e and Fig. 8f, g). Finally, having this highly sensitive and localized calcium sensor, allowed us to follow spontaneous network activity and observe bursts on primary hippocampal neurons using the slow and medium speed sensors (Fig. 8h, i).

**Figure 8.**
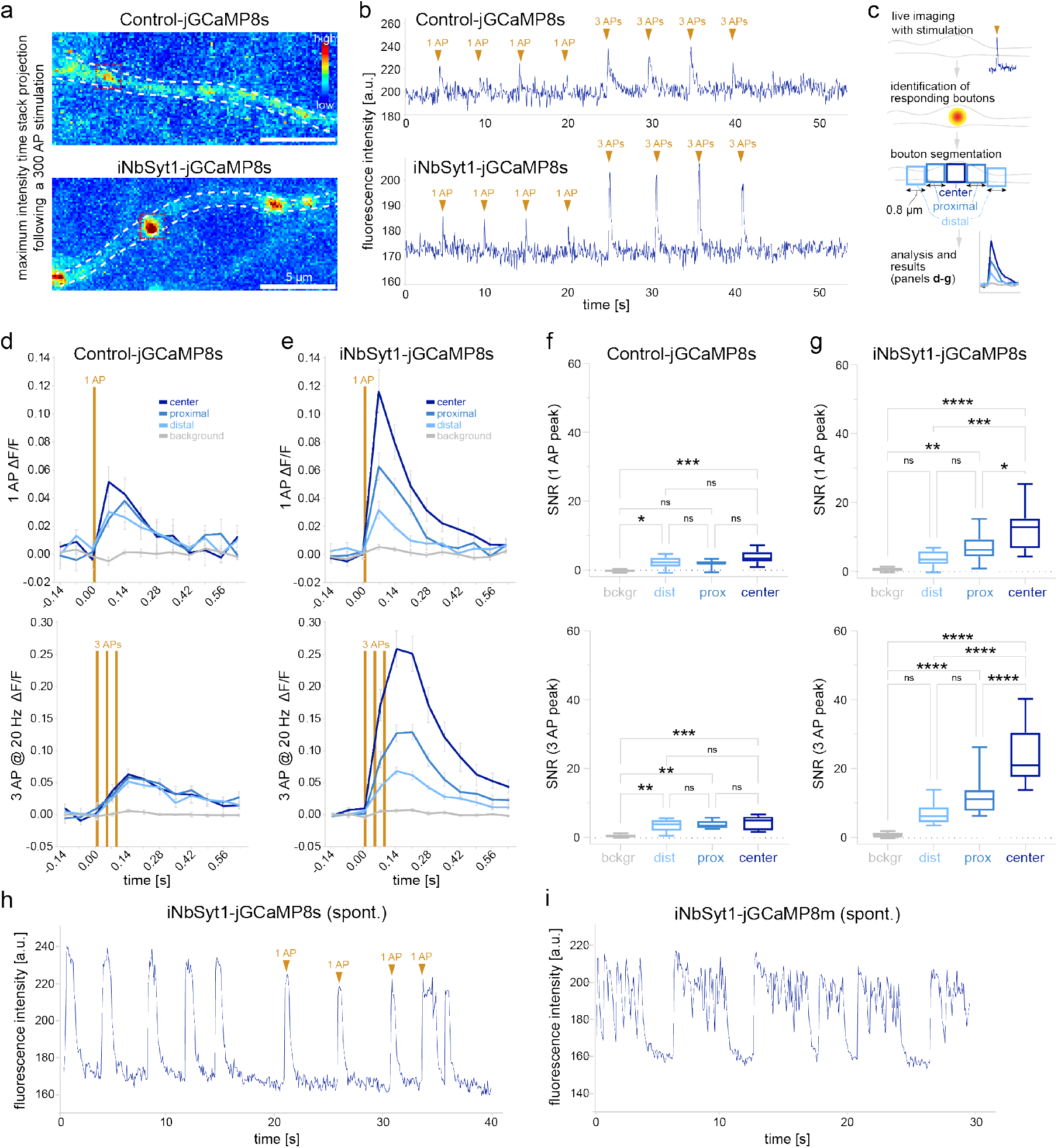
A synaptically localized calcium sensor to precisely detect calcium changes with improved intensity and spatial resolution. (**a**) Expression of the iNbSyt1 fused to jGCaMP8s shows clear presynaptic localization when compared to the control (iNbALFA). Scale bar 5 μm. (**b**) Exemplary traces from single boutons following trains of 1AP or 3AP applied at defined time intervals. Note that in some cases, due to low signal-to-noise ratio (SNR), it is difficult to observe 1AP responses with the control sensor. (**c**) Scheme summarizing the analysis performed in panels d-g. Briefly, at the end of each experiment, a strong 300 AP stimulation was applied to identify responding boutons. For the analysis, the central area of each bouton was considered (center) together with the proximal or the distal synaptic regions, defined as small ROIs of 0.8 μm adjacent to the center as schematized. (**d-e**) Signal expressed as ΔF/F for the different synaptic regions for either the control or for the iNbSyt1-jGCaMP8s following either 1AP or 3APs. Note that the response at the center of the bouton is higher than in the periphery for the iNbSyt1-jGCaMP8s (panel e). N = 80 corresponding to measures obtained for 20 boutons for each condition for each stimulation pattern (coming from 3 independent coverslips). (**f-g**) SNR analyses of the data from panels d and e. The average noise was calculated for each trace at rest and the SNR was measured for a background region and for the regions as schematized in panel c. The data are expressed as box plots where the whiskers represent the 5-95 percentiles. For statistical significance, one way ANOVA followed by Tukey’s multiple comparison tests was used. *P*-values: * = *p* <0.05; ** = *p* < 0.01; *** = *p* < 0.001 and **** = *p* < 0.0001. (**h-i**) exemplary traces of spontaneous activity (without CNQX & D-AP5) measured respectively for the iNbSyt1-jGCaMP8s (h) or iNbSyt1-jGCaMP8m (i). In panel (h) also exogenous stimulation (1AP) is able to trigger network responses. Note that, during bursts, the faster jGCaMP8m version might be appropriate for revealing faster changes in the calcium signal as previously published^19^.

## Discussion

Here we developed and characterized an extremely flexible affinity tool, the NbSyt1, that allows the study of Syt-1 from several angles. We prove the usefulness of this Nb in conventional microscopy, various super-resolution imaging modalities, and as an intrabody in living neurons to follow the molecular physiology of synapses without overexpressing a tagged synaptic protein.

We selected a camelid single-domain antibody binding with excellent specificity and affinity to the C2A calcium-binding domain of Syt-1. The structural data we obtained from the Syt-1-NbSyt1 complex confirms the very high binding affinity (~0.7 nM) and specificity, which is a consequence of polar and hydrophobic interactions, explaining its selectivity for Syt-1. NbSyt1 does not recognize the C2A domain of Syt-2, which shares a very high degree of identity with Syt-1 (77,7% identity on the full protein, and 86,6% on the C2A domain). Importantly, the crystal structure also shows that the binding occurs on the opposite side from the calcium-binding pocket of the C2A domain, which encouraged us to use this nanobody in living neurons as an intrabody, as this region is likely to be not involved in lipid or other protein interactions^10,14^.

The advantages of nanobodies as affinity probes for imaging over classical antibodies have been thoroughly described^24,33,34^. Some of the main benefits are their monovalency (stoichiometric labeling) and their small size allowing to place the fluorophore ~2–4 nm from the target. These features make NbSyt1 ideal for quantitative imaging^35–37^ and high-precision nanoscopy, respectively, where conventional antibodies would place the fluorophores with a linkage error comparable to or sometimes larger than the resolution that different techniques attain^24,38^.

Additionally, the NbSyt1 is functional when expressed in neurons as an intrabody (iNbSyt1). This can be clearly observed *in vivo* as the iNbSyt1 is able to get enriched on presynaptic boutons with practically no effect on SV release physiology (Fig. 6). This feature results in the possibility of specifically labelling synapses relying on the endogenous expression levels of Syt-1 without changing the stoichiometry of synaptic proteins by over-expression or modifying their function or location by fusing them with tags like fluorescent proteins. Therefore, we demonstrated using SpH (Fig. 6) and electrophysiology (Fig.7) strategies that synaptic vesicle machinery appears not to be affected by the presence of iNbSyt1. A not significant tendency of an increased miniature rate when expressed iNbSyt1 could be related to the observations made previously by Courtney et al.^8^.

Finally, we exploited one of its unique features of bringing a sensor directly to the presynaptic compartment, and we fused the iNbSyt1 to a genetically encoded calcium sensor (jCaMP8), demonstrating that it enriches at synapses. This has a simple practical implication as it makes it very convenient for deciding presynaptic regions to image where Syt-1 is enriched. Moreover, having a calcium sensor directly positioned where the calcium should matter the most for SV-release regulation, in close proximity to its sensor, results in the precise and locally-enhanced detection of a single AP with an increased signal-to-noise ratio. Similarly, we used this tool for recording the spontaneous activity of hippocampal neuronal cultures using two chimeras with different speeds in detecting Ca^2+^ variations (respectively the slower iNbSyt1-jCaMP8s and the faster iNbSyt1-jCaMP8m). These two tools will serve the community depending on different experimental needs both for the study of presynaptic calcium and for the role of Syt-1 in different cellular models.

Overall, here we established, engineered and thoroughly characterized a nanobody binding to the C2A domain of Syt-1. This versatile tool shows outstanding features for quantitative and super-resolution microscopy, but it also provides exceptional properties to be used in living neurons and follow or manipulate presynaptic molecular physiology. We expect this bio-tool to be instrumental in understanding Syt-1’s role in SV fusion and neuronal physiology.

## Material and Methods

### Synaptosomes preparations

Rat synaptosomes were enriched as previously described^39^ Briefly, rat brains were homogenized using a glass-Teflon homogenizer in precooled sucrose buffer (320 mM Sucrose, 5mM HEPES, pH 7.4). Centrifugation at 1000 × g for 2 minutes was performed, and the supernatant was further centrifuged at 15,000 × g for 12 minutes. Next, a discontinuous Ficoll density gradient was applied. The fractions at the interface of the 9% Ficoll were pooled and washed in sucrose buffer.

### Immunizations

Two alpacas were immunized with this preparation. The procedure was performed by Preclinics GmbH (Postdam, Germany). Six injections were performed weakly with 500 μg rat synaptosomes (total protein determined by BCA assay). Two weeks after the last immunization a single boost with 500 μg of synaptosomes was performed and 100 ml of blood was taken 3 and 5 days after the boost immunization. PBMCs were isolated using Ficoll gradient and Serum was stored at −80°C. Total RNA was extracted using RNA extraction Qiagen kit (Qiagen).

### Enrichment of IgG2 & IgG3 from plasma

Plasma from the 2 fully immunized animal was enriched in IgG2 and IgG3 following the protocol described by Hamers-Casterman with minimal modifications^40^. The affinity selection steps were done using an Äkta-Prime FPLC system (Cytiva). Between 5 and 10 mL of plasma was diluted in PBS in a one-to-one ratio and filtered through a 0.45 μm pore size syringe filters (Stedim Minisart^®^, Sartorius). The plasma was then injected in HiTrap protein G HP (Cytiva), the flowthrough was collected and injected in HiTrap protein A (Cytiva). The bound IgGs were eluted from the HiTrap protein G column with first 0.15 M NaCl, 0.58% Acetic acid, pH 3.5 to collect mainly IgG3 and a second elution using 0.1 M glycine–HCl, pH 2.7 to collect mainly IgG1. The IgG bound to the HiTrap protein A column were eluted with 0.15 M NaCl, 0.58% Acetic, pH 4.0 to collect the IgG2. To neutralize the pH of the collected fractions, 1 M Tris-HCl pH 9.0 was added and the buffer was exchanged to PBS by injection in HiTrap Desalting Columns (Cytiva). A sample of the collected and desalted fractions were analyzed by denaturing PAGE. IgG2 and IgG3 fractions were pooled together, and residual IgG1 were further removed by incubating the pooled sample with agarose beads conjugated with anti-llama light chain (Capralogics).

### Plasma-ELISA

Purified antigen was immobilized on a 96 well immunosorbent plate (Nunc). All the following steps were done by gentle shaking on an orbital shaker. 30 nmol of purified Syt-1_(197-421)_ was diluted in 200 μL of 100 mM Tris, 150 mM NaCl (pH 8.0) were coated overnight at 4°C. The plate was then washed with PBS and blocked with 5% (w/v) skim Milk in PBS for 3 h at RT. The enriched IgG2-IgG3 were added to the wells in a concentration of 0.5 mg/mL and incubated for 2h at RT. A series of three washes with PBS was performed. The presence of bound IgG2 and IgG3 was revealed with the mouse anti-Camelid antibody coupled to HRP (Preclinics, clone: P17Ig12) diluted 1:2000 in PBS. The antibody was incubated for 2 h at RT, then washed 3 times with PBS. The ELISA was revealed by addition of 100 μL of TMB substrate (ThermoFischer) until the blue color was stable. The reaction was quenched by addition of 100 μL of 2 M sulfuric acid. The absorbance was then read at 430 nm using a plate reader (BioTek).

### Nanobody library generation

Total mRNA was extracted from the PBMC obtained from 2 alpacas using standard RNA extraction kit (Qiagen). Recovered mRNA was retrotranscribed to cDNA by using Supercript IV (ThermoFischer) and the Cal 0001/2 primers as described before^41^. Next, a second PCR was performed to introduce the Gibson cloning overhangs for further insertion in the phagemid. The final PCR product was diluted to 5 ng/μL and some loaded on a 1.5 % Agarose gel to confirming the right size of the PCR product. Fragments were cloned into the phagemid using Gibson assembly. The phagemid backbone used was obtained from the pHen2 plasmid. After Gibson cloning, the obtained construct was purified by PCR purification kit (Qiagen) and the concentration was measured by Nanodrop. The constructs were then electroporated in TG1 bacteria (Biocat). For the transformation, 65 ng of DNA were added to 50 μL of TG1. This process was repeated 20 times. The reactions were left 1 hour at 37°C for initial growth and then pooled in 400 mL of 2YT medium (ThermoFischer) supplemented with antibiotics and grown overnight at 37°C. Next day, bacteria were pelleted and resuspended in 25 mL LB medium (ThermoFischer) and 25 % Glycerol. The Synaptosome-library was aliquoted, snap frozen and stored at −80 °C.

### Phage-Display

The procedure was performed as described in Maidorn et al.^42^ with some modifications. To start the process, a 1 mL the synaptosome nanobody library was diluted in 500 mL of 2YT supplemented with antibiotic and grow at 37°C until OD600 reached ~0.5. Next, ~0.1×10^13^ M13KO7 Helper Phages (NEB) were added to the culture and let the infection for 45 minutes at 37°C. Bacteria were then pelleted and resuspended in 500 mL 2YT medium supplemented with the necessary antibiotics. Infected bacteria were incubated overnight at 30°C to produce the phages. The next day culture supernatant was incubated with 4 % (w/v) PEG-8000 and let on ice for <2 hours to allow phages to precipitate. After several washings steps in PBS, phages were filtered with a 0.45 μm syringe filter (Sartorius). Purified Syt-1_(97-421)_ was conjugated to desthiobiotin-N-Hydrosuccimide Ester (Beryy and Associates). Excess of dt-Biotin-NHS was removed using Nap10 column (Cytiva). Between 1 and 3 nmol of antigen was bound pre-equilibrated Dynabeads MyOne Streptavidin C1 (ThermoFischer) for 1 hour at RT. After binding of Syt-1_(197-421)_ to the beads, the beads were washed 5 times with PBS supplemented with 0.01 % (v/v) Tween (PBS-T). The purified phages were mixed with the beads loaded with Syt-1_(197-421)_ and incubated for 2 hours at RT. Bead were thoroughly washed in PBS-T. The elution of Syt-1_(197-421)_ and bound phages was carried out using 50 mM Biotin in PBS. Eluted phages were then used to reinfect TG1 cells and initiate another cycle of panning (a total of 3 panning rounds were performed). After the last panning round, bacteria were plated in LB agar supplemented with antibiotics. The next day 96 colonies were picked and grew in 96-deep-well plates. A copy of the plate was stored at −80°C after adding 50% glycerol.

### Phage-ELISA

Each clone in the 96-deep-well plate was infected with helper phages and let overnight producing phages. Bacteria are centrifuged down and supernatants full with phages can be use directly. The desthio-biotinilated Syt-1_(97-421)_ was immobilized at 4°C overnight on a flat-bottom Nunc MaxiSorp™ 96-well plate (ThermoFischer). The plate was washed 3 times with PBS-T and blocked using 5% skim Milk and 1% BSA in PBS-T for 2h at RT. Next, 25 μl of phages produced by the picked colonies were incubated on the well with 75 μL 5% skim milk in PBS-T for 1 hour at RT. The unbound phages were washed six times with PBS-T, and bound phages were detected with anti-major coat protein M13-HRP (Santa Cruz, #sc-53004-HRP) diluted 1:1000 in 100 μL PBS-T. The antibody was let for 1 hour at RT and excess was thoroughly washed away with PBS-T. TMB substrate (3,3’,5,5’-tetramethylbenzidine, 1-StepTM Ultra TBM-ELISA, Thermo Scientific) was added to each well and the colorimetric reaction was stopped by addition of 100 μL of 2M sulfuric acid. The absorbance was read at 450 nm throughout the plate (Cytation™ 3: BioTek™ Instruments, Inc.).

### Protein expression and purification

Syt-1 fragments were produced in NEB-Express (NEB), while nanobodies were produced in SHuffle^®^ Express (NEB). Bacteria were grown on terrific broth supplemented with kanamycin at 37°C for Syt-1 and 30°C for nanobodies. When OD600 reached ~2-3, 0.4 mM IPTG was added and temperature was set to 30°C. Induction was allowed for ~16h (overnight). Fully grown cultures were centrifuged and pellet resuspended in cold lysate buffer (LysB: 100 mM HEPES, 500 mM NaCl, 25 mM imidazole, 2.5 mM MgCl_2_, 10% v/v glycerol, 1 mM DTT, pH 8.0) supplemented with DNAse (1:250), lysozyme (1:250) and 1 mM PMSF. After 30 minutes incubation and disruption by sonication, lysate was centrifuged at ~11,000g for 1h at 4°C. Supernatant was incubated with LysB equilibrate Ni^+^ beads (Roche cOmplete Resin) for 1h at RT. Beads were washed with 3 column volumes using LysB buffer, 5 CV with high salt buffer (HSB; 50 mM HEPES, 1.5 M NaCl, 25 mM imidazole, 2.5 mM MgCl_2_, 5% v/v glycerol, 1 mM DTT, pH 7.5). Finally, before the elution, beads were washed in the buffer of choice for the next application. Elution was carried out using self-produced SUMO protease cleaving on column the HisTag-SUMO domain, releasing the protein of interest. Fractions were collected until absorbance a 280 nm drops to baseline. Eluted proteins were initially evaluated on a PAGE.

### Fluorophore conjugation to nanobodies

Purified nanobodies bearing one or two ectopic cysteines (at their C-terminus or N- and C-termini) were reduced for 1 h using 10 mM of tris(2-carboxyethyl)phosphine (TCEP), pH 7. The excess of TCEP was removed using a gravity NAP5 column (Cytiva) previously equilibrated with degasses PBS pH 7.4. Freshly reduced nanobodies were immediately mixed with ~3-5 molar excess of maleimide functionalized fluorophore and incubated for 1h. The excess of dye was removed by using Superdex™ 75 increase 10/300 GL column (Cytiva) on Äkta FPLC system.

### Oligonucleotide conjugation to nanobodies

NbSyt1 was produced having one ectopic cysteine on its C-terminus, while secondary anti-mouse IgG1 nanobody and anti PSD-95 nanobody carrying an ectopic cysteine were obtained from NanoTag Biotechnologies GmbH (#N2005 and #N3705). Nanobodies used for DNA-PAINT imaging were coupled with different docking single-DNA-strands (Biomers GmbH, Ulm, Germany) as described earlier^21,22^. In brief, nanobodies were reduced with 5 mM TCEP, pH: 7, (Sigma-Aldrich, #C4706) for 2 h. After removal of TCEP via 10 kDa molecular weight cut-off (MWCO) Amicon spin filters (Merck, #UFC500324), the reduced nanobodies were coupled using 10-fold excess of maleimide-DBCO crosslinker (Sigma-Aldrich, #760668). The excess of crosslinker was removed using 10 kDa MWCO Amicon spin filters. Azide functionalized DNA-strands (Biomers) were mixed in excess with nanobody-DBCO to promote the azide-alkyne cycloaddition (click) reaction. The excess Azide docking DNA-strands were removed by size exclusion chromatography using an Äkta pure 25 system (Cytiva) equipped with the Superdex^®^ Increase 75 column (Cytiva). The sequences of the docking strands P3 (3’-TTTCTTCATTATTTT-5’) and R4 (3’-ACACACACACACACACACA-5’)^43^,^44^

### Dot blot assays

Target proteins and negative control (BSA) were spotted (~1 μg and ~5 μg respectively) on a nitrocellulose membrane and let them dry at RT. Membranes were incubated with blocking buffer (PBS supplemented with 0.05% (v/v) of Tween20 and 5% milk) under gentle shaking for 2h. After blocking step, blocking buffer was removed and membranes were incubated with NbA51 or NbA91 directly labeled with AbberiorStar635p in 5% milk PBS-Tween20 at a final concentration of 5 nM for 1h. For specificity assay in Supp. Fig 2, one membrane was incubated with 1:100 dilution of polyclonal affinity purified anti-Synaptotagmin 1/2 cytosolic domain (SySy, #105 003) pre-mixed with 3 molar excess of FluoTag^®^-X2 anti-Rabbit IgG conjugated to AbberiorStar635p (NanoTag Biotechnologies, #N2402-Ab635P). Membranes were thoroughly washed with PBS-T and fluorescence signal was detected in Amersham™ Imager 600.

### Cell culture

COS-7 cells were cultured in Dulbecco’s MEM supplemented with 10% FBS, 4 mM L-glutamine, 0.6% penicillin and streptomycin, cells were cultured at 37°C, 5% CO2 in a humified incubator. COS-7 fibroblasts were obtained from the Leibniz Institute DSMZ—German Collection of Microorganisms and Cell Culture (DSMZ Braunschweig, Germany). For immunostainings, cells were plated on poly-L-lysine (PLL)-coated coverslips.

Rat primary hippocampal neuron cultures for imaging were prepared as described before^28^. In brief, the brains of P1-2 rat pups were extracted and placed in cold HBSS (ThermoFisher). The hippocampi were extracted and placed in a solution containing 10 mL DMEM (ThermoFisher), 1.6 mM cysteine, 1 mM CaCl2, 0.5 mM EDTA, 25 units of papain per mL of solution, with CO2 bubbling, at 37°C for 1h. The solution was removed and the hippocampi were incubated in 10% FBS-DMEM, 73 μM albumin for 15 minutes. The hippocampi were triturated using a 10 mL pipette in complete-neurobasal medium (Neurobasal A (ThermoFisher), containing 2% B27 (ThermoFisher) and 1% Glutamax-I (ThermoFisher)). Neurons were plated (12-well plate) on glass coverslips coated with poly-L-lysin-hydrochloride (1 mg/ml, Merck). After 2h, the plating medium was replaced with 1.25 ml complete-neurobasal medium and neurons were incubated for 15 days at 37°C, 5% CO2 in a humidified incubator. Cultures for electrophysiology recordings were prepares according to Goslin and Banker^45^ using embryonic day 18 old rat embryos. Cells were plated in a density of 40.000 cells per 18 mm coverslip, grown in 1 ml of neurobasal medium (NB, Gibco) supplemented with B27 medium. Ara-C (cytosine β-D-arabinofuranoside) at a final concentration of 5 μM was included in the culture medium at DIV 7 to suppress glia proliferation.

### Affinity determination

The affinity of the NbSyt1 was measured by microscale thermophoresis using the device NT.115Pico Monolith (NanoTemper). NbSyt1 was labelled with Alexa647 (as described above) and diluted in MST buffer (NanoTemper) supplemented with 0.05 % Tween. Fluorescent NbSyt1 was incubated with different dilutions of purified Syt-1_(97-421)_ using Premium Coated Capillaries (NanoTemper). For operation of the instrument and evaluation of affinity data, the MO.Control and MO.Affinity Analysis software (NanoTemper) were used.

### Immunofluorescence

For Fig. 1b COS-7 Cells were fixed for 20 minutes using 4% paraformaldehyde (PFA) in PBS at RT. After rinsing short with PBS, remaining aldehyde groups were quenched for 15 minutes using 0.1 M glycine in PBS at RT. Cells were blocked and permeabilized with 3% (w/v) BSA + 0.1% (v/v) Triton X-100 for 40 minutes at RT and gently shaking. Nanobody candidates were applied in PBS supplemented with 1.5% BSA and 0.05% Triton X-100 for 2h at RT with gentle shaking. After staining, several washing steps using PBS were carried out including DAPI short staining. Coverslips were shortly rinsed in distilled water and mounted using Mowiöl (12 ml of 0.2 M Tris buffer, 6 ml distilled water, 6 g glycerol, 2.4 g Mowiol 4-88, Merck Millipore). Samples were imaged directly or within the next 48h, samples were kept at 4°C. Primary hippocampal neuronal cultures (15 DIV) imaged in Fig. 1c and Fig.3 were first fixed with 4% PFA solution for 30 minutes at RT and quenched with 0.1 M glycine (Merck) solution in PBS for 15 minutes at RT. Samples were permeabilized and blocked with PB buffer (1.5% BSA and 0.05% Triton X-100 in PBS at pH 7.4) for 40 minutes at RT. For immunostaining, mouse monoclonal anti-Synaptotagmin 1 (#105011, SySy) recognized by conventional secondary Ab conjugated to Cy3 (1:500, #115-005-166, JIR; for Fig. 1) or for Fig. 3a and g, the anti-α-Tubulin primary Ab ((#302211, SySy) was premixed with FluoTag-X2 anti mouse IgG1 coupled to Atto488 (#N2002, NanoTag Biotechnologies) initially in 20 μl of PBS and after 30 minutes at RT, 980μl of PBS was added to bring the concentration of Ab and nanobody to 7 and 15 nM respectively. Into this pre-mixture tube, 25 nM of FluoTag-X2 anti-PSD95 coupled to AbberiorStar580 (#N3702, NanoTag Biotechnologies) and 25 nM NbSyt1 coupled to AbberiorStar635p were added. Samples were incubated for 1 hour at RT under slow orbital stirring. Neurons were washed 3x with PBS and rinsed once with high salt PBS (0.5 M NaCl). Samples were finally rinsed in water and mounted on glass slide using Mowiol. For DNA-PAINT imaging (Fig. 3b, c, d, e and f) immunostaining was performed using ~50 nM of the NbSyt1 functionalized with the docking DNA strand P3 coupled to Atto488 on its 3’-end (NbSyt1-P3-Atto488). Simultaneously, post-synapses were labelled with 50 nM of FluoTag-X2 anti-PSD95 (#N3702, NanoTag Biotechnologies) functionalized with the docking DNA strand R4 on its 3’-end (NbPSD95-R4). After immunostaining, samples were post-fixed with 4% PFA for 15 minutes, quenched with 0.1 M Glycine and stored in PBS in 4 °C until imaged.

### Crystallization and structure determination

NbSyt1 was incubated with the cytosolic C2A_(140-265)_ domain of Synaptotagmin 1 at 1:1 molar ratio and concentrated to 65 mg/ml using 3000 Da cut-off spin columns at 4°C, prior to setting sitting-drop crystallization plates. Crystals of the NbSyt1-Synaptotagmin 1 complex suitable for data collection were found in the G8 condition from the JCSG+ screen (Molecular Dimensions), containing 0.15 M Di-Malic acid pH 7.0, 20% PEG 3350 at 1:1 volume ratio with the protein solution. Crystals were frozen and data collection was performed at the BioMAX beamline, MAX IV (Lund, Sweden)^46^. The data were processed using XDS^47^, molecular replacement was performed with Phaser^48^ using structures 5T0R^49^ and 6I2G^32^ as search models. The structure was built using Coot^50^ and refined in Phenix^51^. Data collection and refinement statistics are shown in Supp. Table 1.

### Live intrabody imaging and electrical stimulation in hippocampal neurons

Images were taken by an inverted Nikon Ti epifluorescence microscope (Nikon Corporation, Japan) equipped with a Plan Apochromat 60×, 1.4 NA oil immersion objective, an IXON X3897 Andor camera, an Oko Touch Environmental control and NIS-Elements software.

#### Live imaging of intrabodies

Rat hippocampal neurons (DIV ~11) were transduced with an adeno-associated virus (AAV) containing the sequence for expression of directly fluorescently labelled (mScarlet or mNeonGreen) intrabodies (iNbSyt1-ALFA / iNbALFA-3xFLAG) and imaged live 3-4 days post-infection. Transduced neurons were placed onto a custom-built live-imaging chamber and imaged in Tyrode’s solution (124 mM NaCl, 5 mM KCl, 30 mM glucose, 25 mM Hepes, 2 mM CaCl2 and 1 mM MgCl_2_ at pH 7.4) at ~0.2 frames per second (fps).

#### SynaptopHluorin (SpH)

Rat hippocampal neurons (DIV ~11) were infected with an AAV holding a bicistronic plasmid for expression of the respective intrabody and SpH. Live imaging experiments were performed between day 3-5 post-infection. Transduced neurons were placed onto a custom-built live-imaging chamber and imaged in Tyrode’s solution supplemented with 10 μM cyanquixaline (CNQX, Tocris Bioscience, Cambridge, UK) and 50 μM 2-amino-5-phosphonopentanoic acid (D-AP5; Tocris Bioscience, Cambridge, UK) to avoid spontaneous network activation. Neurons underwent field stimulation at 30s (for 3s at 20Hz; i.e., 60 AP) and for 90s (600 AP) while imaged at a rate of 1 fps. After electrical stimulation, 50 mM NH4Cl was applied to evaluate the total SpH pool^31^.

#### Calcium imaging

Rat hippocampal neurons (DIV ~10) were infected with an AAV leading to expression of the respective intrabody fused to the Ca^2+^-sensor jGCaMP8s/m^19^. Live imaging experiments were performed between day 4-6 post-infection. Transduced neurons were placed onto a custom-built live-imaging chamber and imaged in Tyrode’s solution. For the assessment of spontaneous network activity, neurons were imaged for 60s in the absence of blockers. For responses following field stimulation, the Tyrode’s solution was supplemented with 10 μM CNQX and 50 μM D-AP5 to avoid spontaneous network activation. Neurons underwent field stimulation at 20Hz using one field stimulation (one action potential, 1AP) or 3APs. Images were taken at a rate of 15 fps.

The analysis of images with these sensors was performed using Fiji/imageJ^52^. For synaptic endocytosis, briefly, based on the total content of SVs (signal uncovered with NH4Cl treatment), regions of interest were set and fluorescence intensity measured and averaged for each time point per image. These averaged time courses then were normalized to fluorescent intensities directly preceding the respective stimulation trigger, bleach corrected and peak aligned. For jGCaMP8s analyses, the responding boutons were identified at the end of the movie applying 300 APs. The ΔF/F was calculated measuring the basal fluorescence, averaging the fluorescence for at least 5 time points before each stimulation. This basal initial fluorescence (F) was used to calculate the ΔF (corresponding to the fluorescence at each time point from which the initial fluorescence F was subtracted), divided by F. This ΔF/F is to account for possible small differences in the expression of the sensors across different cells.

### Confocal & STED microscopy

Images from mounted samples in Mowiol and expanded gels in a custom-made chamber were acquired using STED Expert line microscope (Abberior Instruments). The microscopy setup was comprised of an IX83 inverted microscope (Olympus) equipped with UPLSAPO 100x 1.4 NA oil immersion objective (Olympus). In addition, 488 nm, 561 nm and 640 nm lasers were used for confocal imaging. High-resolved images were obtained from the same setup using the 595 nm and 775 nm pulsed STED depletion lasers in combination. Images were analyzed in Fiji/ImageJ (v. 1.53o).

### Structured Illumination Microscopy (SIM)

Imaging was performed on the Elyra 7 (Zeiss) with Lattice SIM^2^ microscope equipped with a Plan-Apochromat 63x 1.4 NA oil immersion objective (Zeiss). Highly resolved 3D 3-coloured images were acquired on 80 × 80 μm field of view after overlapping 40 z-stacks. For that, glass slides of immunostained neurons mounted in Mowiol were exited with 642nm, 561nm and 488nm laser wavelengths. Image reconstruction and analysis took place in the ZEN software (Zeiss), after acquiring several frames from each z-stack.

### Exchange-PAINT and FL-PAINT imaging and data analysis

Both Exchange-PAINT and FL-PAINT measurements were performed on a custom-built confocal setup^53^. The imaging chamber was fixed with clips to the microscope’s stage and a PDMS layer was used as top cover. Prior to the super-resolution imaging, an area of 80 μm × 80 μm was scanned and then a suitable region for imaging was selected based on the NbSyt1-Atto488 signal. Typically, 20k scan images with virtual pixel size of 100 nm and pixel dwelling time of 2.5 μs were recorded. The total acquisition time was around 45 minutes for a 20 μm × 20 μm scan region. The laser power was adjusted according to the sample brightness. In FL-PAINT experiments, a mix of imager DNA strands R4-Atto550 (5’-TGTGTGT-Atto550-3’), which labeled the NbPSD95 and P3-Cy3B (5’-GTAATGAAGA-Cy3B-3’), which labeled NbSyt1, were used at a final concentration of 0.1 nM each in PBS buffer including 500 mM NaCl. For Exchange-PAINT imaging, the solutions exchange in experimental chamber has been realized using custom-built microfluidics system ^21^. First, R4 imager DNA strand was diluted in PBS buffer including 500 mM NaCl to the final imaging concentration of 0.1 nM. After imaging the first target, the imager was washed away from the chamber by flushing the chamber with PBS buffer. Then, the imaging buffer with the next imager was introduced into the chamber and the next imaging round has been performed. In Exchange-PAINT and FL-PAINT data analysis, we chose a time binning of three scanned frames, corresponding to the time bin of 0.3 s, while the scan region was 20 μm × 20 μm. The detection of emitter candidates and its sub-pixel localization were performed using a cross-correlation algorithm and pixel-integrated Gaussian MLE fitting^22^. Blinking events detected in a single frame were discarded. Out-of-focus localizations with point spread function (PSF) width of more than 180 nm or/and a number of detected photons in a single event smaller than 100 were rejected. For lifetime determination, we discarded the first 0.1 ns after the maximum of the TCSPC histograms, and tail-fitted the remained curve of a histogram with a mono-exponential function using a maximum likelihood estimator (MLE). Lifetime values in the range from 0.5 to 5.0 ns were included in the analysis. Subsequently, a drift correction was applied, and finally DNA PAINT images were reconstructed. The following average localization precision values were calculated: 11.9 nm for Figure 3c, 13.6 and 8.4 nm for Figure 3f (P3 and R4 imagers). For targets crosstalk estimation in the FL-PAINT imaging in Fig 3c, we fitted the lifetime histograms with two-Gaussian function and determined lifetime threshold values manually to separate between the two different targets. Then, we calculated the total crosstalk as the total number of localizations in-between thresholds (wrong attribution), divided by the total number of localizations beyond thresholds (correct attribution), as shown in Supp. Figure 4.

### Expansion Microscopy

Primary hippocampal neurons were washed once with pre-warmed PBS (37°C) and fixed using 4% PFA solution for 10 minutes at 4°C and then for 30 minutes at RT. Unreacted aldehydes were quenched with 100 mM NH4Cl in PBS for 20 minutes at RT. Neurons were washed three times with permeabilization and blocking (PB) buffer (2% BSA, 0.1% Triton-X 100 in PBS). For immunolabeling, NbSyt1 equipped with anchoring cassettes and AbberiorStar635p together with nanobody anti-PSD95 (FluoTag-X2 anti-PSD95, #N3705; NanoTag Biotechnologies) conjugated to sulfo-AlexaFluor488 were used at 20 nM in PB buffer for 1h at RT. Excessive probes were washed with PB buffer and then samples were postfixed using 4% PFA in PBS for 10 min at RT followed by quenching with 100 mM NH4Cl in PBS for 15 minutes at RT. Next, samples were undergone 10x Expansion^26^ with some modifications. In Brief: Acryloyl-X (#A20770, ThermoFisher) stock solution (5 mg) was diluted in DMSO reaching the concentration of 10 mg/ml (pH: 8) followed by further dilution to 0.3 mg/ml in PBS and 500 μl was added on the cells. Samples were incubated overnight at RT in the dark and washed with PBS before subjected to gelation. Fresh monomer solution (MS) was prepared by adding 1.335g N,N-Dimethylacrylamide (DMAA, #274135, Sigma) stock solution and 0.32g sodium acrylate (SA, #408220, Sigma) in 2.85 ml of ddH2O. Potassium persulfate solution (KPS) was freshly prepared at a stock of 36 mg/ml and added at 3.6 mg/ml in degassed ice-cold MS, protected from light. For the gelation reaction 4 μl TEMED (#T7024, Sigma) was added to 1 ml of KPS/MS solution, vortexed and a droplet (90 μl) was immediately spotted on parafilm. Thereafter, samples were placed with cells down on the droplet and left for ~24h at ~23°C until the gel was fully polymerized. For Homogenization, gel was detached from the parafilm, transferred into a glass petri dish and was incubated with the disruption buffer (5% Triton-X and 1% SDS in 100 mM TRIS, pH 8.0) for 2 hours with disruption buffer exchange at 30 min intervals. Next, samples were autoclaved for 20 minutes at 121°C in disruption buffer and gels were transferred into 120mm square petri dish filled with ddH2O and expanded by exchanging the water every 30-60 minutes 4-5 times. Expanded gels were left overnight in water and placed in custom-made chamber for imaging.

### Molecular biology

Sequences for intrabody expression were synthesized (ThermoFischer) fused to a C-terminal ALFA-tag and cloned into an AAV backbone^54^, which additionally contained an expression cassette for Synpatobrevin2-pHluorin (SynaptopHluorin or SpH). Both genes, the intrabody and the SpH are under the control of a hSyn1-promoter. For the constructs used in calcium imaging, the first hSyn1-promoter together with the Vamp2-pHluorin cassette was removed. Subsequently, the desired sequence for jGCaMP8s/m was cloned in frame to NbSyt1-ALFA or NbALFA-3xFLAG by restriction sites for AgeI and AscI. For live imaging of the intrabodies NbSyt1 (iNbSyt1) and NbALFA (iNbALFA), the nanobody sequences were fused to mScarlet or mNeonGreen. Fluorescent proteins were amplified with the following primers (forward: TCCACCGGTACCATGGTGAGCAAGGGCGAGG and reverse: ATTGGCGCGCCTCTTGTACAGCTCGTCCATGCC) to introduce the restriction sites AgeI and AscI and cloned into the AAV backbone containing the iNbSyt1-ALFA or iNbALFA-3xFLAG.

### Adeno-associated viruses (AAVs)

The AAVs were generated as previously described^54,55^. AAVs were produced in HEK293T cells by co-transfection of pHelper plasmids (pFΔ6, pRV1, p21) and pAAV target plasmid in a 4:1:1:2 molar ratio by use of Lipofectamine 2000 (ThermoFisher) according to company protocol. 72 h post transfection, cells were harvested and lysed by 3 cycles of thawing and freezing followed by treatment with benzoase nuclease (Millipore). After pelleting down cell debris (16.9 g/rcf, 30 minutes at 4°C), supernatants were filtered, aliquoted and snap frozen in liquid nitrogen, before viruses were stored at −80°C until used.

### Electrophysiology

All experimental procedures were carried out in accordance with the EU Council Directive 2010/63/EU and were approved by the local Committee for Ethics and Animal Research (Landesverwaltungsamt Sachsen-Anhalt, Germany). Electrophysiology was performed on primary hippocampal cells infected with AAV encoding for the intrabody and SpH from DIV14-18 under visual control using oblique illumination on a Slicecope Pro2000 (Scientifica, Uckfield, East Sussex, UK). Prior to recordings, expression was verified pHluorin fluorescence. Whole-cell voltage-clamp recordings were performed using a MultiClamp 700B amplifier, filtered at 8 kHz and digitized at 20 kHz using a Digidata 1550A digitizer (Molecular Devices). Extracellular Tyrode solution, adjusted to pH 7.4 and 300 mOsm, contained: 150 mM of NaCl, 4 mM of KCl, 1.25 mM of MgCl_2_.6H2O, 10 mM of glucose, 10 mM of HEPES, 2 mM of Ca2+, and 1 μM of tetrodotoxin (TTX). Intracellular pipette solution for voltage clamp recordings, adjusted to pH 7.3 and 285 mOsm, contained: 115 mM of Cs-methanesulphonate, 10 mM of CsCl, 5 mM of NaCl, 10 mM of HEPES, 20 mM of TEA.Cl hydrate, 4 mM of MgATP, 0.3 mM of GTP, 0.6 mM of EGTA, and 10 mM of QX314. Whole-cell recordings were carried out at −60 mV (reversal potential of GABAA receptors) for mEPSCs and 0 mV (reversal potential of AMPA and NMDA receptors) for mIPSCs. For statistical analysis (Fig. 7), two-tailed unpaired Mann-Whitney U test was performed. It is a nonparametric test in which the sample medians are compared based on the shape of distribution between the two independent groups. To correct significance level, a Bonferroni correction was then employed. The resulting box plots display the median and the first and third quartile with whiskers extending to 1.5 times the interquartile range above and below the upper and lower quartiles, respectively (Q1 - 1.5 * IQR or Q3 + 1.5 * IQR).

## Acknowledgments

FO was supported by Deutsche Forschungsgemeinschaft (DFG) through the SFB1286 (project Z04). EFF was supported by a Schram Stiftung (T0287/35359/2020) and a Deutsche Forschungsgemeinschaft (DFG) grant (FO 1342/1-3). EFF also acknowledges the support of the Collaborative Research Center 1286 on Quantitative Synaptologie (CRC/SFB1286), Göttingen, Germany. PS and MMC acknowledge the MAX IV Laboratory for time on Beamline BioMAX under Proposal 20190233. Research conducted at MAX IV, a Swedish national user facility, is supported by the Swedish Research Council under contract 2018-07152, the Swedish Governmental Agency for Innovation Systems under contract 2018-04969 and Formas under contract 2019-02496. This work was supported by grants from the Novo Nordisk Foundation (NNF20OC0064789), the Swedish Research Council (2022-03681) and the Swedish Cancer Society (20 1287 PjF) to PS. MP and HH were supported by the Leibniz “Best Minds” program and the Center for Behavioral Brain Sciences (CBBS). We thank Silvio Rizzoli for critical reading of the manuscript and Christina Zeising, Nicole Hartelt and Christina Patzelt for the excellent technical help.

## Author contributions

KQZVR, NM, RR and SSI designed, performed and analyses most of the experiments. LA helped with calcium imaging and its analysis, AMR and KAS planed and performed ExM. MM initiated the phage-display screenings, JH produced and conjugated proteins. MMC and PS planned, performed and analyzed the structural biology work. HH and MP planned, performed and analyzed electrophysiology data. NO and RT helped with imaging and analysis of Exchange PAINT and FL-PAINT. EFF, designed and guided experiments, performed analysis and wrote the manuscript. FO conceived and lead the project, designed performed and guided experiments, analyzed data, and wrote the manuscript. All authors contribute with writing.

