## Supplementary material for "A versatile synaptotagmin-1 nanobody provides perturbation-free live synaptic imaging and low linkage-error in super-resolution microscopy": Supp. Fig.

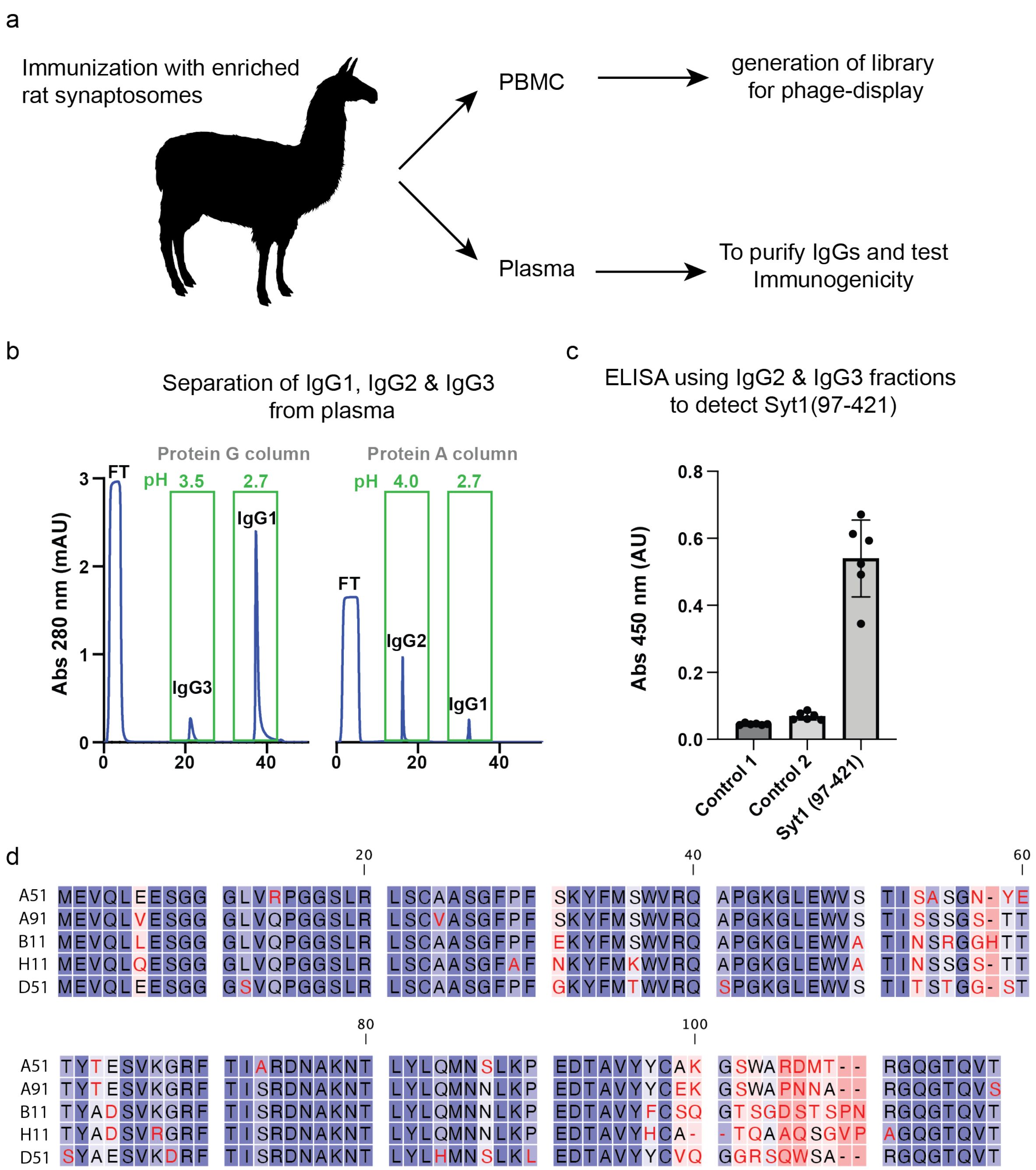


**Supplementary Figure 1.** Overview of immunization and discovery strategy**.** (**a**) Immunization was performed using synaptosomes enriched from rat central nervous system. After immunization campaign, peripheral blood lymphocytes (PBMCs) and plasma were separated from 100 ml of blood. PBMCs were further use to generate the nanobody library, while plasma was further process to separate the different antibodies isotypes. (**b**) Example of the chromatographic procedure to enrich for IgG2 and IgG3 from plasma using a combination of Protein G and Protein A columns with elutions at different pH (FT: flow through of material not bound to column). (**c**) rSyt-1$_{(97\text{-}421)}$ was immobilized on an ELISA plate and revealed using enriched IgG2 and IgG3 as obtained in (b). Negative control 1 correspond to wells without rSyt1 immobilized and control 2 are wells without IgG2/3. For both controls we use the same amount and times of the detection system (anti alpaca antibody conjugated to Horse radish peroxidase, followed by Turbo TMB substrate). (**d**) Sequences of the 5 clones showing the higher signal in ELISA when rSyt-1 was immobilized.

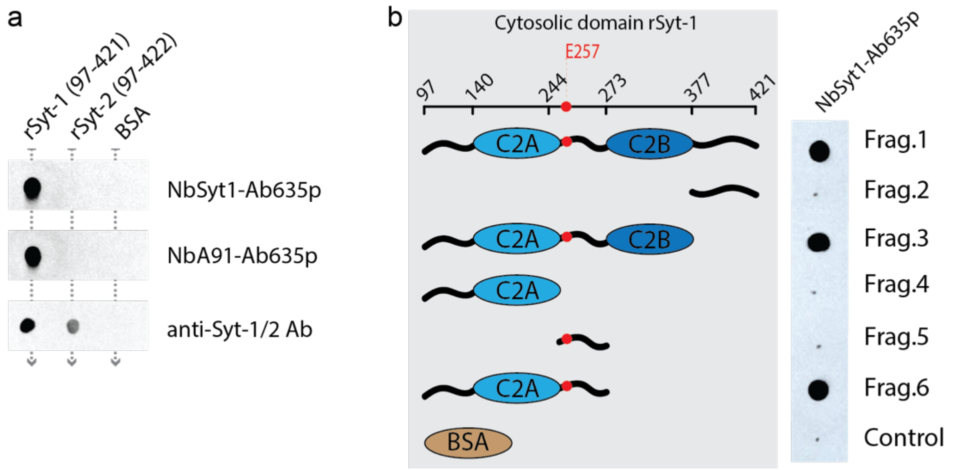


**Supplementary Figure 2.** Specific for Syt-1, epitope mapping. (**a**) Equivalent amounts of purified rat Syt-1$_{(97\text{-}421)}$ and rat Syt-2$_{(97\text{-}422)}$ were spotted on nitrocellulose membranes, as control 5 µg of bovine serum albumin (BSA) was also spotted on the membrane. After blocking, equivalent membranes were incubated with nanobodies directly labeled with AbberiorStar635p (NbSyt1-Ab635p, NbA91-Ab635p) or using conventional primary, wash secondary antibody (rabbit polyclonal primary recognizing both Syt1/2 (SySy, Cat# 105 003), revealed by secondary antibodies conjugated with AbberiorStar635p). (**b**) different truncated fragment from the cytoplasmic domain of rat Synaptotagmin 1 (rSyt-1) were spotted in equivalent amount on a nitrocellulose membrane, together with BSA as control (Fragment 1 (Frag.1): 97-421; Frag.2: 377-421; Frag.3: 97-377; Frag.4: 97-244; Frag.5: 244-273; Frag.6: 97-273). NbSyt1-Ab635p was used to reveal the recognized fragments.

**Supp. Table 1**. Data collection and refinement statistics.

| | |
|---|---|
| **Data collection** | |
| Space group | P 21 21 21 |
| Cell dimensions | |
| *a, b, c* (Å) | 52.41, 65.75, 66.91 |
| α, β, γ (°) | 90.0, 90.0, 90.0 |
| Resolution (Å) | 1.68 |
| Rsym or Rmerge | 0.087 (3.822) |
| *I/σI* | 11.85 (0.66) |
| Completeness (%) | 97.8 (96.1) |
| Multiplicity | 8.9 (9.2) |
| **Refinement** | |
| Resolution (Å) | 1.68 |
| No. reflections | 26393 (1246) |
| $R_{work}/R_{free}$ | 0.19/0.23 |
| **No. atoms** | 2051 |
| Protein | 1967 |
| Ligand/ion | 37 |
| Water | 47 |
| ***B*-factors** | 37.69 |
| Protein | 37.44 |
| Ligand/ion | 49.76 |
| Water | 38.64 |
| **R.m.s. deviations** | |
| Bond lengths (Å) | 0.006 |
| Bond angles (°) | 0.89 |

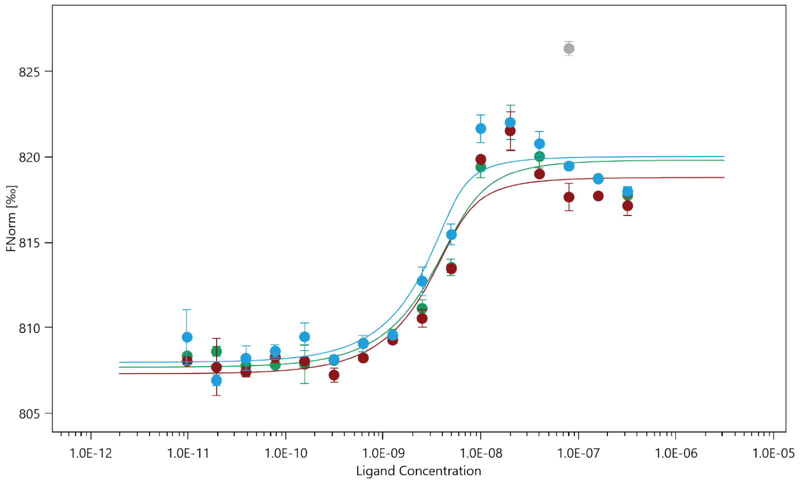


### Dataset Overview ●

| Name: | First Sample | Second Sample | Third Sample |
|---|---|---|---|
| Graph Color: | ● | ● | ● |
| Target Name: | Nb A51 | Nb A51 | Nb A51 |
| Target Concentration: | 5 nM | 5 nM | 5 nM |
| Ligand Name: | Synaptotagmin1 | Synaptotagmin1 | Synaptotagmin1 |
| Ligand Concentration: | 320 nM to 0.00977 nM | 320 nM to 0.00977 nM | 320 nM to 0.00977 nM |
| n: | 2 | 2 | 2 |
| Comments: | | | |
| Excitation Power: | 5% | 5% | 5% |
| MST Power: | 60% | 60% | 60% |
| Temperature: | 22.0°C | 22.3°C | 21.9°C |
| **Kd:** | **1.0883E-09** | **7.0506E-10** | **4.1627E-10** |
| Kd Confidence: | ± 7.7984E-10 | ± 6.5832E-10 | ± 4.6011E-10 |
| Response Amplitude: | 12.115881 | 11.476413 | 12.027773 |
| TargetConc: | 5E-09[Fixed] | 5E-09[Fixed] | 5E-09[Fixed] |
| Unbound: | 807.69 | 807.31 | 807.98 |
| Bound: | 819.81 | 818.79 | 820.01 |
| Std. Error of Regression: | 1.3749205 | 1.4772043 | 1.4553421 |
| Reduced $\chi^2$: | 24.55617 | 171.57472 | 570.00506 |
| Signal to Noise: | 9.5181117 | 8.3452543 | 8.8775696 |

**Supplementary Figure 3.** Affinity determination using microscale thermophoresis (nanoTemper). Three independent measurements of directly labeled NbSyt1-Alexa647 mixed at different concentrations with rSyt-1$_{(97\text{-}421)}$.

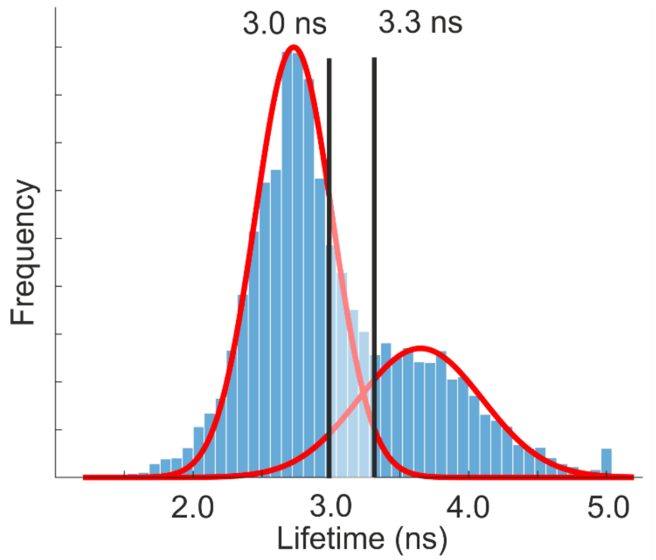


**Supplementary Figure 4:** Target crosstalk evaluation in FL-PAINT. Lifetime histogram (blue), the fitted Gaussian function (red), and the selected lifetime threshold values (black lines) of image shown in Figure 3b. The crosstalk is calculated based on lifetime threshold values. To improve separation between the targets, localizations in-between two lifetime threshold values were excluded (the region between the peaks marked in light blue color). In this case the two-target total crosstalk is 3.8%.
